# Endothelial cells are a key target of IFN-g during response to combined PD-1/CTLA-4 ICB treatment in a mouse model of bladder cancer

**DOI:** 10.1101/2023.03.28.534561

**Authors:** Sharon L. Freshour, Timothy H.P. Chen, Bryan Fisk, Haolin Shen, Matthew Mosior, Zachary L. Skidmore, Catrina Fronick, Jennifer K. Bolzenius, Obi L. Griffith, Vivek K. Arora, Malachi Griffith

## Abstract

To explore mechanisms of response to combined PD-1/CTLA-4 immune checkpoint blockade (ICB) treatment in individual cell types, we generated scRNA-seq using a mouse model of invasive urothelial carcinoma with three conditions: untreated tumor, treated tumor, and tumor treated after CD4+ T cell depletion. After classifying tumor cells based on detection of somatic variants and assigning non-tumor cell types using SingleR, we performed differential expression analysis, overrepresentation analysis, and gene set enrichment analysis (GSEA) within each cell type. GSEA revealed that endothelial cells were enriched for upregulated IFN-g response genes when comparing treated cells to both untreated cells and cells treated after CD4+ T cell depletion. Functional analysis showed that knocking out *IFNgR1* in endothelial cells inhibited treatment response. Together, these results indicated that IFN-g signaling in endothelial cells is a key mediator of ICB induced anti-tumor activity.

## Introduction

Bladder cancer is a common malignancy worldwide (6th most common among men and 17th most common among women) and accounts for over 500,000 new cancer diagnoses and 200,000 cancer-related deaths per year.^1^ While over 95% of bladder cancer cases are classified as urothelial carcinomas, they encompass a range of molecular subtypes, which are primarily distinguished by differential expression of differentiation markers and may predict for response to specific treatments.^2, 3^ Initial diagnosis stages can be broadly grouped into non-muscle invasive (NMIBC), muscle invasive (MIBC), and metastatic disease. About 75% of cases are initially diagnosed as NMIBC, 20% as MIBC, and the remaining 5% as metastatic. Depending on the initial degree of invasiveness and metastasis, 5-year survival rates can range from 96% to 6%.^4^

Standard treatment recommendations likewise depend on the initial degree of invasiveness as well as the risk stratification of recurrence and progression. NMIBC is typically treated with a transurethral resection followed by either chemotherapy, Bacillus Calmette-Guerin immunotherapy, or radical cystectomy in high risk cases.^2^ MIBC is typically treated with neoadjuvant chemotherapy followed by radical cystectomy and, in some cases, adjuvant immunotherapy. Previous research has suggested that response to treatment may differ by subtype. For example, basal/squamous bladder cancers may have better response to neoadjuvant chemotherapy than luminal-infiltrated tumors.^3^ While neoadjuvant chemotherapy has historically been used most commonly, clinical trials looking at the use of neoadjuvant immune checkpoint blockade (ICB) treatments have shown promise as well.^5, 6^

Currently, there are several ICB treatments approved by the FDA for treatment of bladder cancer, all of which are either PD-1 or PD-L1 inhibitors.^7, 8^ Initially, these treatments were approved specifically for treatment of advanced disease, targeting patients who were ineligible for cisplatin treatment.^9^ Over time, ICB use has become more widespread and has been applied across the range of bladder cancer stages from NMIBC to metastatic disease.^8, 10, 11^ While ICB therapy shows great promise for treatment of bladder cancer, there are still many patients who do not receive benefit from ICB treatment. Thus, there remains a need to improve treatment methods, determine which patients will respond well to treatment, understand mechanisms of response to treatment, and identify potential predictors of response.^5^

Clinical trials examining the benefit of PD-1/PD-L1 inhibitors in urothelial carcinoma have found that high IFN-g expression is associated with treatment response, suggesting that IFN-g signatures could serve as a predictor of response.^12^ Additionally, treatment response has been associated with high expression of *CXCL9* and *CXCL10*, two IFN-g induced chemokines that have been associated with increased T cell infiltration in multiple tumor types.^13, 14^ However, these trials did not fully explore how or where IFN-g may be acting to help induce or improve treatment response. Clinical trials have also looked at improving treatment response by combining PD-1/PD-L1 inhibitor treatments with CTLA-4 inhibitor treatments. These trials have shown greater response rates compared to PD-1/PD-L1 monotherapy.^15–18^ Nevertheless, the challenges of identifying ideal patients for treatment as well as identifying mechanisms and predictors of response remain.

To study mechanisms of response to combined PD-1/CTLA-4 ICB treatment, we used a murine muscle-invasive urothelial carcinoma cell line generated by exposing mice to 4-hydroxybutyl(butyl)nitrosamine (BBN), which caused them to develop areas of invasive disease. These tumor bearing bladders were then resected and used to propagate an organoid cell line, MCB6C.^19^ Previous analysis showed that MCB6C is responsive to ICB treatments and achieves the best treatment response with combined PD-1/CTLA-4 ICB treatment. Additionally, this previous work showed that treatment response was dependent on CD4+ T cells, consistent with research showing that CD4 T cells may be the primary mediators of anti-tumor activity in human bladder cancer.^20^ Analysis of the MCB6C model also showed that ICB treatment led to expansion of IFN-g producing CD4+ T cells with a Th-1 like phenotype. Neutralizing IFN-g in the tumor negated the anti-tumor activity of combined treatment, indicating that IFN-g was a key mediator of response. Surprisingly, this research showed that knocking out *IFNgR1* in the tumor cells themselves did not affect treatment response, suggesting that IFN-g was mediating treatment response through non-tumoral cells in the tumor microenvironment (TME).^19^

To better understand mechanisms of treatment response in this model, we performed single cell RNA sequencing (scRNA-seq) on MCB6C tumors isolated from mice under three conditions: untreated tumor, tumor treated with combined PD-1/CTLA-4 ICB treatment, and tumor treated with combined ICB treatment after CD4+ T cell depletion (Figure 1A). For each condition in each replicate, tumors from three mice were resected and pooled to generate single cell suspensions for 10x 5’ gene expression sequencing as well as B cell and T cell receptor sequencing (Figure 1B, Methods). This sequencing was performed for five biological replicates. In addition to scRNA-seq, whole genome and exome sequencing of the tumor cell line were performed, along with matched normal whole genome and exome sequencing of a tail sample (Figure 1C, Methods).

**Figure 1:**
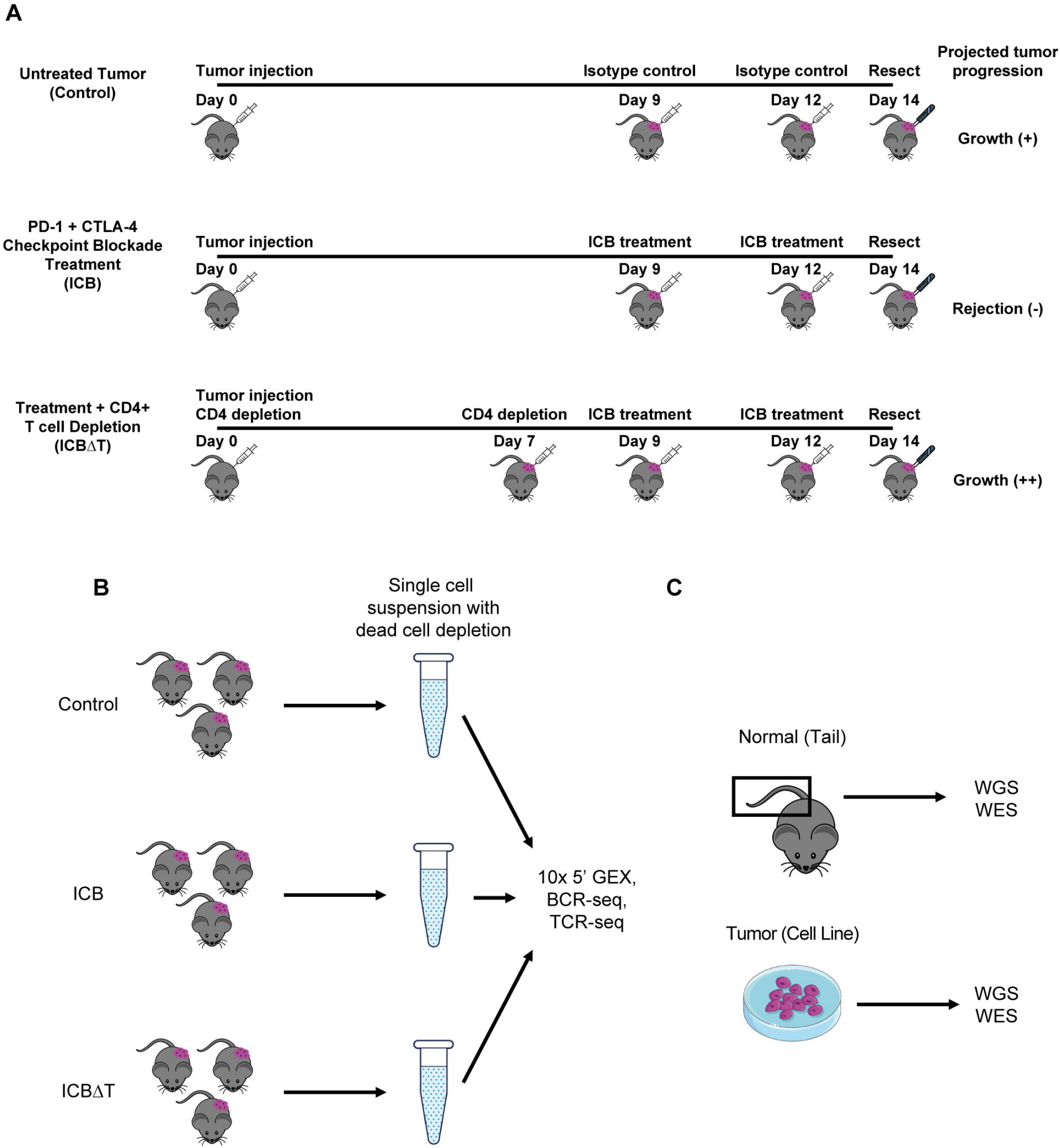
Experimental design for single cell RNA and bulk DNA sequencing. (A) Timelines for generating tumor samples for individual mice for each condition. (B) Workflow for generating single cell suspensions for single cell RNA sequencing for one of five biological replicates sequenced. For each condition in each replicate, tumors from three individual mice were pooled into one suspension and used to create 10x libraries for single cell gene expression (GEX), B cell receptor (BCR) and T cell receptor (TCR) sequencing. (C) Sources for normal and tumor bulk DNA sequencing. DNA was isolated from a normal mouse tail sample and an MCB6C tumor cell line sample for whole genome sequencing (WGS) and whole exome sequencing (WES). ICB = combined PD-1/CTLA-4 immune checkpoint blockade treatment, ICBΔT = combined PD-1/CTLA-4 immune checkpoint blockade treatment received after CD4+ T cell depletion, Isotype control = rat IgG2a and mouse IgG2b.

## Results

### Bulk DNA sequencing shows that the MCB6C cell line is clonal with a high mutation burden, normal ploidy, and a stable genome

Bulk whole genome sequencing (WGS) of the tumor cell line generated over one billion paired reads, 88% of which produced high quality alignments (i.e. had a mapping score of Q20 or greater). Bulk WGS of the normal tail sample produced over 1.1 billion reads, with approximately 91% of reads having high quality alignments. Bulk whole exome sequencing (WES) of the tumor cell line produced over 55,000,000 reads with over 90% of reads having high quality alignments, while WES of the tail sample produced over 77,000,000 reads with over 90% of reads having high quality alignments (Supplemental Table 1).

After alignment, we performed somatic variant calling with the WES data and identified 16,449 possible somatic variants, including 16,315 single nucleotide variants (SNVs) and 134 small insertions or deletions (indels), before filtering. These variants were then filtered using several metrics, including total coverage, variant allele frequency (VAF), and consensus across callers (Methods). 10,427 variants remained after filtering, of which 10,407 were SNVs and 20 were indels, showing that the MCB6C cell line has a high SNV burden (approximately 4.17 mutations per Mb). Once we identified this high-quality set of likely somatic variants, we checked the clonality of the cell line by plotting the distribution of the VAFs of all SNVs (Figure 2A, Supplemental Table 2). This analysis showed a normal distribution with a median VAF of 48.26%, indicating that the MCB6C tumor cell line is highly clonal.

**Figure 2.**
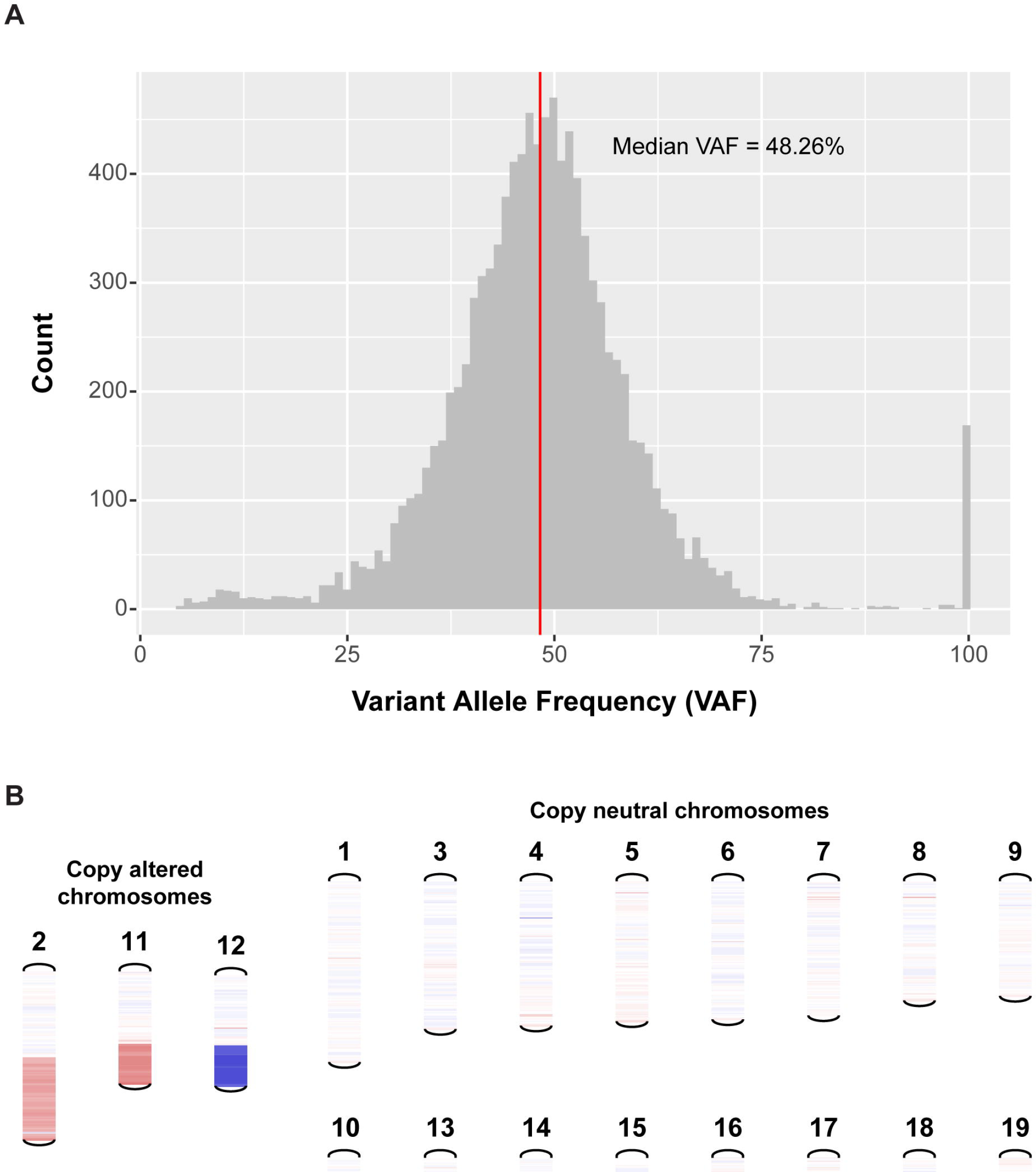
Bulk DNA sequencing shows that the MCB6C cell line is clonal with a high mutation burden, normal ploidy, and a stable genome. (A) Distribution of variant allele frequencies (VAFs) for somatic SNVs detected using matched tumor and normal whole exome sequencing of the MCB6C cell line. Red line indicates overall median VAF (48.26%). Of the 168 total variants appearing at VAF of 100%, 27.4% were found within a region of loss on chromosome 12 and 70.8% were found on chromosome X. (B) Visualization of copy number variants detected using matched tumor and normal whole genome sequencing of the MCB6C cell line. Three chromosomes exhibited copy number variation: chr2 (partial copy gain, i.e., log2 copy ratio of approx. 0.32, of ∼85 Mb), chr11 (single copy gain, i.e., log2 copy ratio of approx. 0.5, of ∼36 Mb), and chr12 (single copy loss, i.e., log2 copy ratio of approx. -0.5, of ∼42 Mb). See also supplemental table 2.

Looking at individual somatic mutations, we confirmed three driver mutations (*Kras* G12D, *Trp53* T122K, and *Kdm6a* H1146Y) for the MCB6C cell line, which were previously reported from analysis of bulk whole transcriptome sequencing (RNA-seq).^19^ Along with these three mutations, we identified 31 additional mutations across 20 previously reported driver genes in human bladder cancer, including a second *Trp53* mutation (a splice donor variant) and a second missense *Kdm6a* mutation (Supplemental Table 2, Methods).^21^ This set also included mutations in *Atm* (S1884T), *Fat1* (two missense, one stop-gained mutations), *Kmt2a* (H1067Q), and *Kmt2c* (one splice region mutation) which have each been shown to harbor mutations in over 10% of bladder cancers, although none of the specific mutations identified appear to have been previously reported in bladder cancer.^3^ In addition to the stop-gained and splice region mutations identified in *Atm* and *Kmt2c*, respectively, two additional stop-gained mutations (one in *Birc6* and one in *Rnf213*) and three additional splice region variants (one in *Birc6*, one in *Brca2*, and one in *Sf3b1*) were identified. Finally, a mutation in *Sf3b1* (E873K), which was identified as a possible driver of a similar mouse urothelial carcinoma cell line, but was not previously detected in MCB6C using RNA-seq, was detected using WES.^19^

In addition to calling SNVs and indels, we called copy number variants using the WGS data (Methods). These results indicated that the MCB6C cell line has a relatively stable genome with only a few copy number events consisting of copy gains on chromosomes 2 and 11 and copy loss on chromosome 12 (Figure 2B, Supplemental Table 2). Together, these results indicated that the MCB6C cell line has high SNV burden and low CNV burden. Previous research has shown that metastatic urothelial carcinoma patients with high SNV/low CNV tumor profiles may benefit more from ICB therapy. While high SNV/low CNV status has been associated with greater chance of response, the utility of SNV and CNV status is still being evaluated as a possible predictor of treatment response in bladder cancer.^22^

### scRNA-seq was generated for over 64,000 cells, with over 59,000 cells passing filtering

10x Genomics 5’ single cell gene expression sequencing (scRNA-seq) of fifteen samples, five per each condition, was performed, generating 8,266,616,441 reads across 64,049 cells. The average number of reads captured per cell in each sample ranged from 82,775 reads to 348,813 reads, with sequencing saturation estimates ranging from 87.0% to 97.1%. For all samples except the treated condition in the fourth replicate (which had 18.4% detection), the fraction of reads detected per cell for each sample was 89.9% or greater. The median number of genes detected per cell for each sample ranged from 380 to 1,864, with the total number of unique genes detected per sample ranging from 14,688 to 20,825 (Supplemental Table 1). In addition to gene expression sequencing, 10x Genomics V(D)J sequencing of B cell receptor (BCR) and T cell receptor (TCR) sequences was performed on all samples, with 14 out of 15 BCR samples successfully run through CellRanger’s V(D)J pipeline and all TCR samples successfully run (Supplemental Table 1).

Before analyzing the scRNA-seq data, we aggregated all three conditions for each replicate and performed basic filtering on each of the five replicates to remove cells that appeared to be low quality based on mitochondrial gene expression per cell, detected gene count per cell, and/or total UMI count per cell. Briefly, cells expressing high percentages of mitochondrial genes, cells with low gene counts, and cells with high UMI counts were removed (Figures 3A-C, Supplemental Table 3, Methods). Ultimately, 4,708 cells across all fifteen samples were removed, with 59,341 cells remaining.

**Figure 3.**
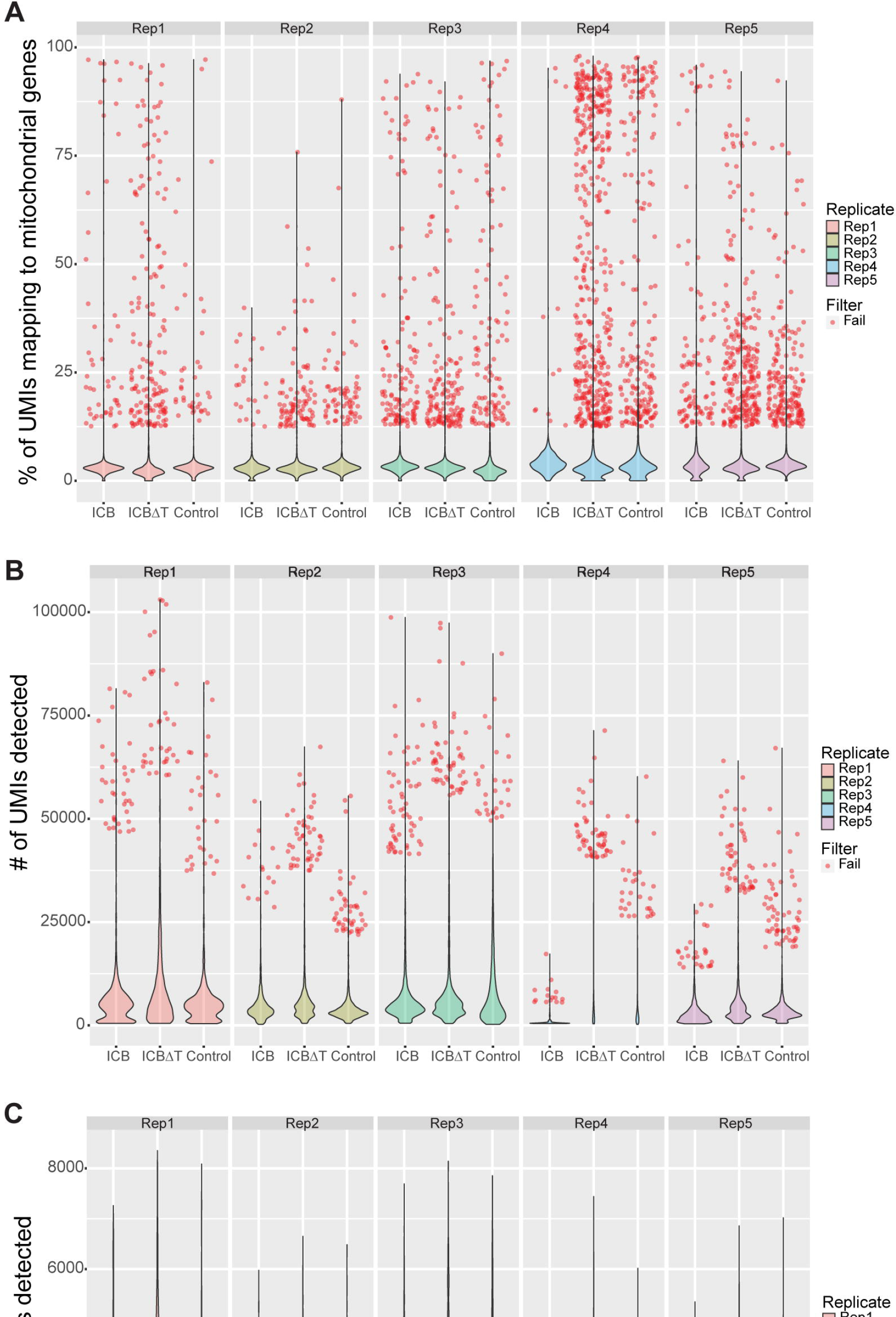
scRNA-seq was generated for over 64,000 cells, with over 59,000 cells passing filtering. (A) Violin plots showing the distribution of the percentage of unique molecular identifiers (UMIs) mapping to mitochondrial genes per cell, for all 64,049 unfiltered cells, split by condition and replicate. Across all replicates and conditions, 2,058 cells failed filtering based on the percentage of mitochondrial gene expression per cell and each cell is plotted individually as a red dot. (B) Violin plots showing the distribution of the total UMI count detected per cell, for all 64,049 unfiltered cells, split by condition and replicate. Across all replicates and conditions, 584 cells failed filtering based on the total UMI count per cell and each cell is plotted individually as a red dot. (C) Violin plots showing the distribution of the total count of unique genes detected per cell, for all 64,049 unfiltered cells, split by condition and replicate. Across all replicates and conditions, 3,132 cells failed filtering based on the total count of unique genes detected per cell and each cell is plotted individually as a red dot. In total, 4,708 cells failed filtering across all three filtering criteria. Note that cells can fail filtering based on more than one criterion and will be plotted as red dots for each criterion failed. ICB = combined PD-1/CTLA-4 immune checkpoint blockade treatment, ICBΔT = combined PD-1/CTLA-4 immune checkpoint blockade treatment received after CD4+ T cell depletion. See also supplemental table 3.

### scRNA-seq allows identification of lymphocyte, myeloid, and stromal cell populations in the tumor microenvironment

After completing basic filtering of cells, we used SingleR with the ImmGen dataset to assign fine label cell types to all remaining cells from each replicate.^23–25^ We then further filtered the set of remaining cells, removing all cells marked as “pruned” by SingleR. “Pruned cells” are those cells that have received poor-quality cell type assignments, potentially because of underlying poor quality of the cell itself. Once we removed all pruned cells, we were left with 57,818 cells total across all conditions and all replicates, which we aggregated into a single gene-barcode matrix for downstream analysis. Before beginning any additional analysis, we also filtered out lowly expressed genes. For a gene to pass filtering, we required the gene to be detected in two or more cells in each replicate, with a supporting UMI count of at least two in each cell. Ultimately, after filtering genes based on these criteria, we were left with 11,398 genes.

Next, we performed manual curation of SingleR’s fine label cell type assignments to group fine labels of the same broad cell type and to identify subtypes within certain broad cell types, e.g., to identify naive CD4 and CD8 T cells within the broader CD4 and CD8 T cell populations (Methods). We also confirmed the general accuracy of the cell type assignments. First, for several cell types, we picked one reported marker for each cell type (e.g. *Cd79a* for B cells, *Epcam* for epithelial cells, and *Col3a1* for fibroblasts) and compared the expression of each marker in the cell type expected to express it (based on the SingleR cell type assignment) versus all other cell types (Figures 4B-E, Supplemental Table 4). These plots confirmed that the expected cell types generally showed more common and higher expression of their markers than non-expected cell types. Additionally, cells identified as expressing B cell receptor (BCR) sequences (i.e. cells that are likely B cells) or T cell receptor (TCR) sequences (i.e. cells that are likely T or NK cells) were compared to cells labeled as either a B or T/NK cells, respectively, according to their gene expression signatures. These results showed that approximately 92.8% of cells identified as expressing BCR sequences were labeled as B cells by SingleR and 98.9% of cells identified as expressing TCR sequences were labeled as some type of T or NK cell, i.e., CD4, CD8, NK, NKT, Tgd, or Treg cells (Figures 4F-G, Supplemental Table 4).

**Figure 4.**
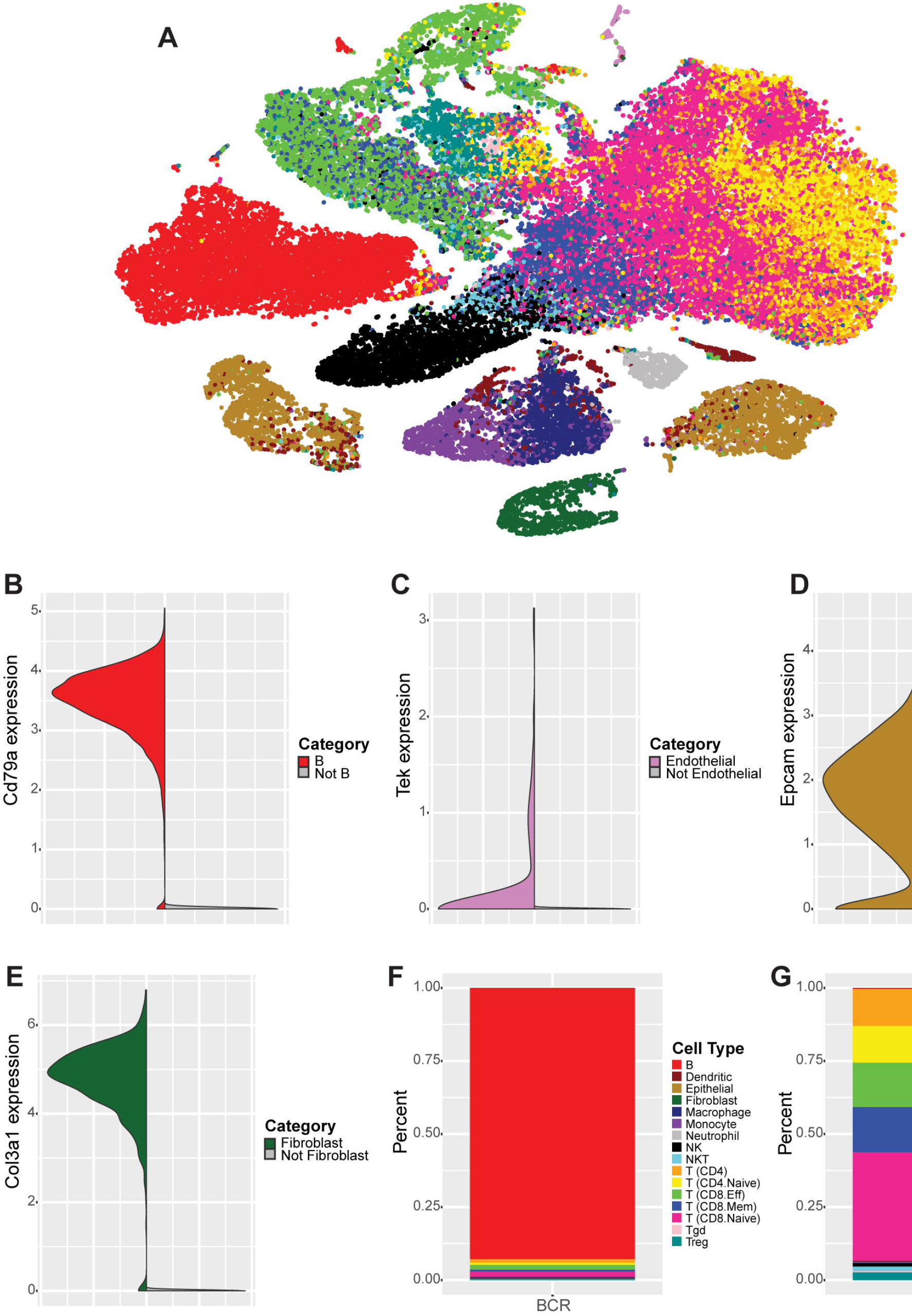
scRNA-seq allows identification of lymphocyte, myeloid, and stromal cell populations in the tumor microenvironment. (A) tSNE clustering projection of the aggregated data set containing 57,818 cells, across all replicates and conditions, that passed filtering and were not “pruned” by SingleR. Cells are colored by manually curated SingleR cell types. (B) Split violin plot showing expression of *Cd79a* in cells labeled as B cells (red, left) versus all other cell types (gray, right). (C) Split violin plot showing expression of *Tek* in cells labeled as endothelial cells (light purple, left) versus all other cell types (gray, right). (D) Split violin plot showing expression of *Epcam* in cells labeled as epithelial cells (gold, left) versus all other cell types (gray, right). (E) Split violin plot showing expression of *Col3a1* in cells labeled as fibroblasts (dark green, left) versus all other cell types (gray, right). (F) Stacked bar plot showing the proportion of cell types assigned to cells identified as expressing B cell receptor (BCR) transcripts by CellRanger’s V(D)J pipeline. (G) Stacked bar plot showing the proportion of cell types assigned to cells identified as expressing T cell receptor (TCR) transcripts by CellRanger’s V(D)J pipeline. See also supplemental table 4.

Once we had confirmed that SingleR’s cell typing was performing as expected, we then generated a tSNE clustering projection for the aggregated data set and colored cells by their manually curated cell type labels (Figure 4A, Supplemental Table 4, Methods). This clustering revealed that although the T lymphocyte populations did not always form distinct clusters on the tSNE projection, within the shared clusters there was often internal structure skewed towards either higher density of CD4 or CD8 T cell types. Additionally, within the CD8 T cell populations, there was clear separation of naive and non-naive populations (i.e. effector and memory CD8 T cells). Similarly, monocytes, macrophages, and some dendritic cells had overlapping clustering with internal separation of the monocyte and macrophage populations.

While the vast majority of B lymphocyte cells clustered separately from other cell types and formed one cluster, we observed a small population of B cells with a distinct expression signature from the main B cell cluster. Based on co-expression of *Cd79a* and *Jchain* in many of these cells, we determined this population likely represented a population of plasma cells (Supplemental Figures 1A-B). Likewise, epithelial cells clustered separately from other cell types, but showed evidence of two distinct populations of similar size within the broader epithelial population (discussed extensively below). Endothelial cells, fibroblasts, and neutrophils largely clustered in single clusters, separate from other cell types.

### Somatic variation can be used to identify tumor cell populations with high confidence

Since we expected tumor tissue to be epithelial, we expected that the epithelial populations identified by SingleR would correspond to the tumor cell populations. To verify this expectation, we classified cells as tumor or non-tumor based on the presence or absence of somatic mutations as follows.

With the 10,427 somatic variants identified from WES, we used VarTrix to detect supporting reads for the reference and alternate alleles at each variant position in each individual cell in the aggregated data set (Figure 5A). To identify a high confidence set of variant-containing cells, we required a cell to have at least two variant positions with greater than 20X total coverage, greater than five reads supporting the alternate allele, and a variant allele frequency (VAF) over 10%. Using these criteria, we classified 4,628 cells as somatic variant-containing cells.

**Figure 5.**
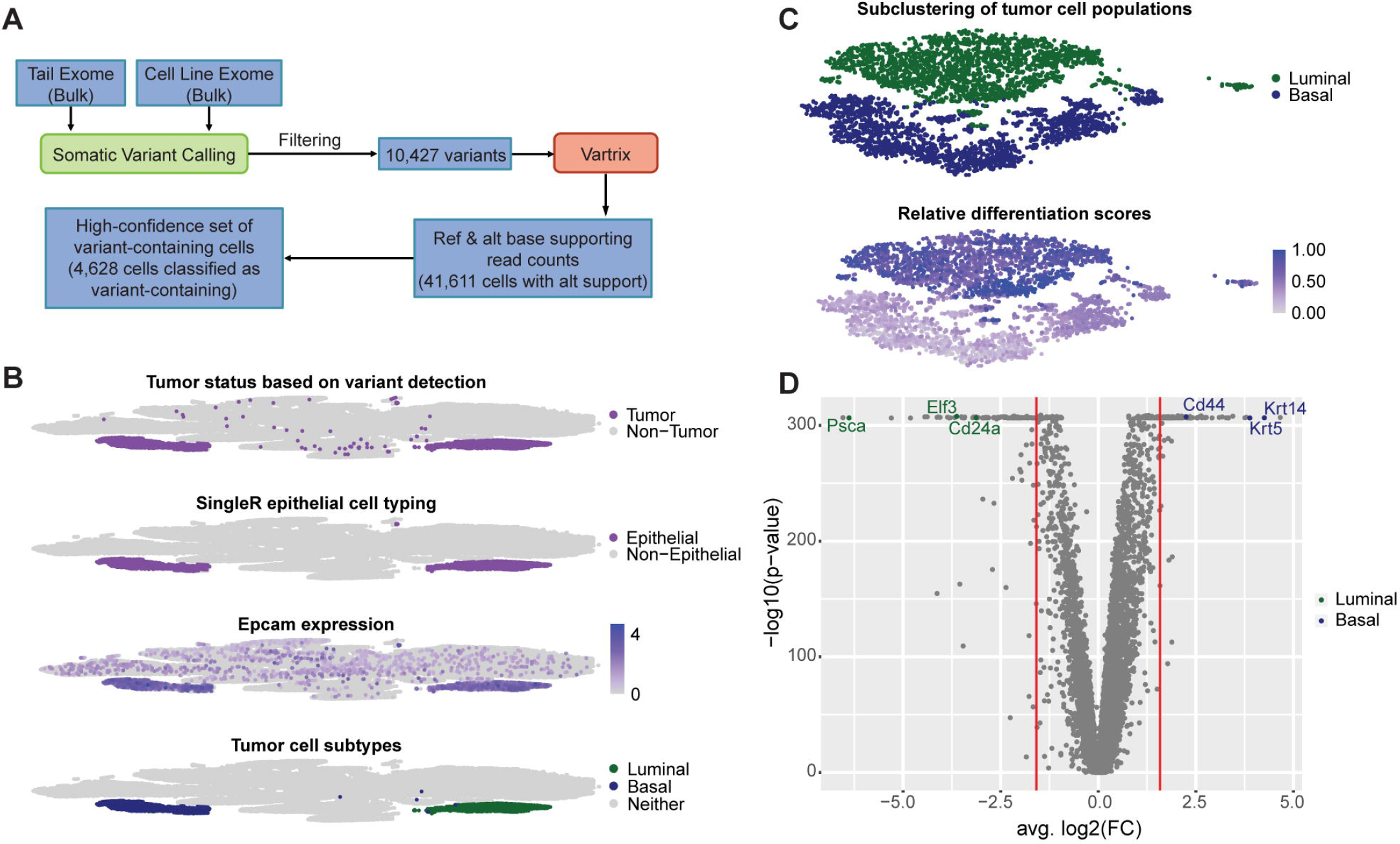
Somatic variation can be used to identify tumor cell populations, which include distinct basal-like and luminal-like subpopulations. (A) Workflow for detecting somatic variation in scRNA-seq data to identify variant-containing cells. (B) tSNE clustering projections showing the classification of tumor cells based on variant detection, epithelial cell typing from SingleR, *Epcam* expression, and subtyping for basal-like and luminal-like tumor cells based on *Krt14* and *Psca* marker expression, respectively. (C) tSNE clustering projections of tumor cell populations showing subtype assignments and differentiation scores (1 - CytoTRACE scores). Differentiation scores indicate the relative differentiation states of each cell within the full tumor cell population. Differentiation scores close to 1.00 indicate cells are relatively more differentiated. Differentiation scores close to 0.00 indicate cells are relatively less differentiated. (D) Volcano plot for differential expression results generated using Seurat’s FindMarkers function to compare the basal-like tumor cell population to the luminal-like tumor cell population. Genes marked in green have been reported in literature as being associated with luminal tumors. Genes marked in blue have been reported in literature as being associated with basal tumors. See also supplemental figure 2 and supplemental tables 5 and 6.

These variant-containing (or variant-positive) cells largely formed two distinct clusters on the tSNE projection, which heavily overlapped the two clusters identified as epithelial clusters using SingleR’s cell type labels (Figure 5B, Supplemental Table 5). Since we expected the tumor tissue to be epithelial tissue, this extensive overlap appeared to confirm that variant-positive status could be used to identify tumor cells with high confidence. Additionally, we compared the overlap of variant-positive cells, cells that were assigned as epithelial cells by SingleR, and cells that were expressing Epcam, a marker of epithelial tissue. While the two variant-positive, epithelial-typed clusters showed high, widespread expression of Epcam as expected, there was also evidence of Epcam expression across numerous other clusters (Figure 5B). These results indicated that variant status can be used to distinguish epithelial tumor cells from possible normal epithelial cells (i.e. cells that are Epcam-positive, but variant-negative).

### Tumor cell populations form two distinct clusters corresponding to basal and luminal subtypes

After confirming which cells and clusters corresponded to tumor cell populations, we investigated why tumor cells formed two distinct clusters. Based on expression of known markers *Krt5* and *Psca*, we provisionally identified one group as a basal-like subpopulation and the other as a luminal-like subpopulation, respectively (Figure 5B, Supplemental Table 5). To further confirm whether these populations represented distinct basal-like and luminal-like tumor subpopulations, we separated the tumor clusters from the rest of the aggregated data set and reclustered them (Methods). The tSNE clustering projection again revealed distinct clustering of each of the two subtypes (Figure 5C). We then assigned relative differentiation scores to each cell and performed differential expression analysis comparing the basal-like cluster, containing all three conditions, to the luminal-like cluster, also containing all three conditions (Methods). The relative differentiation scores revealed that the luminal-like cells largely corresponded to the most highly differentiated cells, while the basal-like cells appeared to form two groups of cells - one which corresponded to the least differentiated cells and one which corresponded to slightly more differentiated, but still relatively lowly differentiated cells (Figure 5C, Supplemental Table 6). These results are consistent with previous literature that has reported that luminal bladder cancers display characteristics of greater differentiation than basal bladder cancers.^26^

After generating differential expression analysis results for comparing the full basal population to the full luminal population, we looked for evidence of reported basal and luminal bladder cancer markers among the most highly differentially expressed genes (Methods). Looking at downregulated genes (i.e. genes that were downregulated in the basal-like population compared to the luminal-like population or, alternatively, genes that were upregulated in the luminal-like population compared to the basal-like population), we found *Elf3* and *Cd24a*, two previously reported markers of luminal bladder cancer, among the most downregulated genes (Figure 5D, Supplemental Table 6).^26, 27^ Looking at upregulated genes in the basal-like population, we found *Cd44* and *Krt14*, two previously reported markers of basal bladder cancer, among the most upregulated genes.^3, 28^ We also confirmed that *Krt5* and *Psca*, the markers used to initially identify the basal-like and luminal-like populations, were among the most up- and downregulated genes, respectively (Figure 5D, Supplemental Table 6). Ultimately, these results showed that there was strong evidence of two distinct molecular subtypes (basal-like and luminal-like) within the tumor cell populations.

Furthermore, overrepresentation analysis of highly differentially expressed genes identified by comparing basal ICB treated cells to luminal ICB treated cells suggested the two subtypes may respond differently to treatment (Methods). In particular, the set of highly differentially expressed genes was enriched for genes relating to IFN-g response (*B2m*, *Gbp2*, *H2-D1*, *Ifitm1*, *Parp14*, *Psme2*, *Rtp4*, *Tmem140*, *Wars*) and IFN-a response (*B2m*, *Cxcl9*, *Gbp8*, *H2-D1*, *Icam1*, *Nfkbia*, *Parp14*, *Psme2*, *Rtp4*, *Wars*), which were primarily downregulated in basal-like ICB treated cells compared to luminal-like ICB treated cells, suggesting that luminal tumor cells may have stronger IFN-g and IFN-a related responses to treatment than basal tumor cells (Supplemental Table 6, Supplemental Figures 2A-B). While this identification of tumor subtypes and differential responses highlights the power of single cell sequencing to detect heterogeneity at a greater level than bulk sequencing, this heterogeneity within the tumor population may not impact overall treatment response in this model since previous research has shown that treatment response is not dependent on IFN-g activity in the tumor itself.^19^

### Overrepresentation and gene set enrichment analysis identify IFN-g response as a commonly perturbed gene set across immune and tumor cell types

After assigning cell types and subtypes, where appropriate, to all cells, we explored how each individual cell type was responding to treatment. To do this, we performed differential expression analysis comparing each possible pair of conditions within each cell type (Methods). We then used the results of these differential expression analyses to perform overrepresentation analysis and gene set enrichment analysis (GSEA).

Overrepresentation analysis revealed that MSigDB’s hallmark IFN-g response gene set was one of the top three most commonly overrepresented gene sets across cell types and comparisons (Figure 6A, Supplemental Table 7, Supplemental Figures 3A-C). The other two most commonly overrepresented gene sets were allograft rejection, which appeared to be reflective of a generalized immune response, and *Tnfa* signaling via *Nfkb*. Given that prior research suggested that IFN-g within the TME may be an important mediator of treatment response, we chose to explore the IFN-g response gene set further.^19^

**Figure 6.**
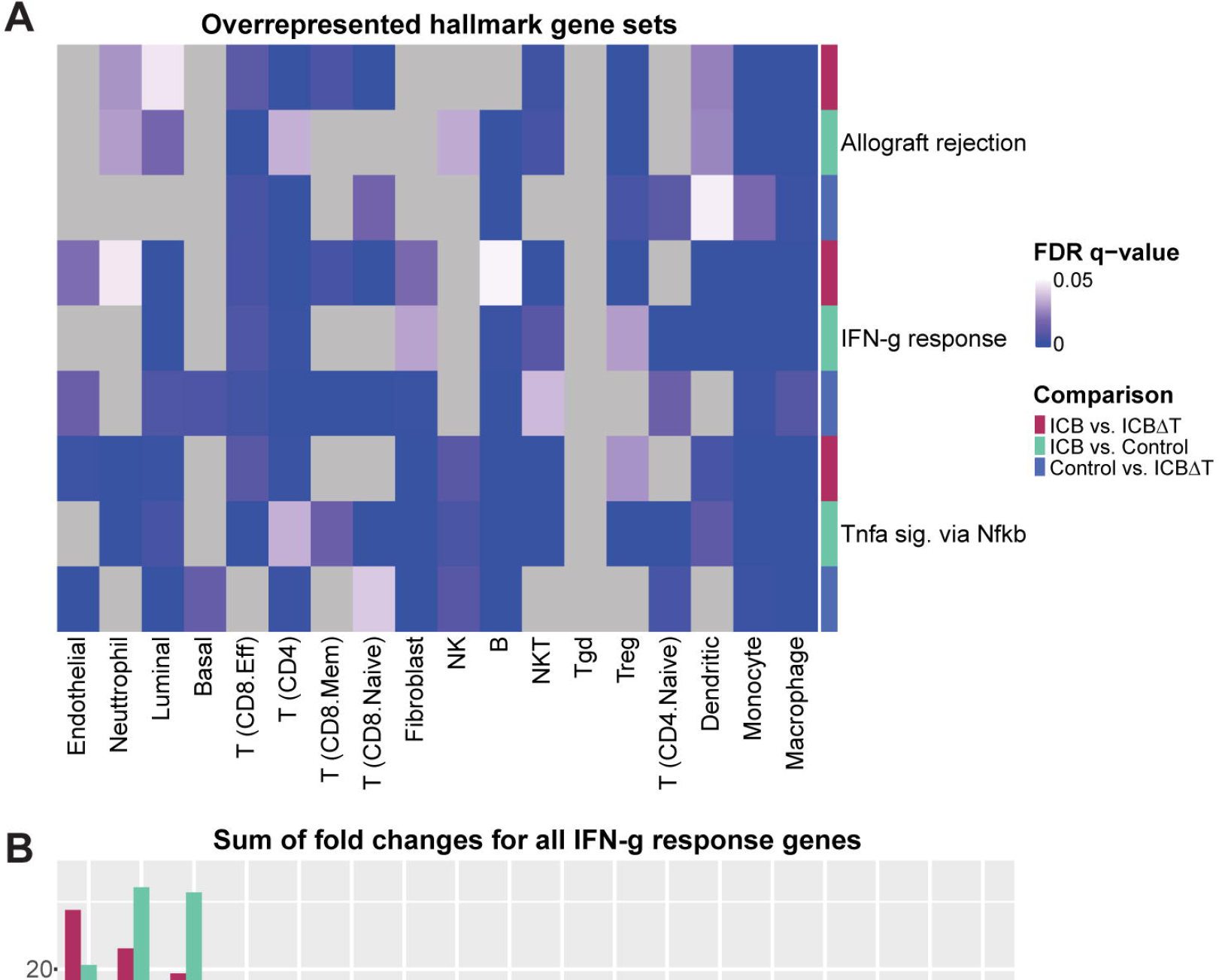
Overrepresentation and gene set enrichment analysis identify IFN-g response as a commonly perturbed gene set across immune and tumor cell types. (A) False Discovery Rate (FDR) q-values for the top three most commonly overrepresented hallmark gene sets for each pairwise comparison of conditions in each cell type. (B) Sum of fold changes for hallmark IFN-g response genes for each pairwise comparison of conditions in each cell type. Positive values indicate that IFN-g genes skew towards upregulation in the first condition of a given comparison. Negative values indicate that IFN-g genes skew towards downregulation in the first condition of a given comparison. (C) Normalized enrichment scores (NES) for the hallmark IFN-g response gene set for each pairwise comparison in each cell type. Positive NES values indicate enrichment of upregulated genes. Negative NES values indicate enrichment of downregulated genes. ICB = combined PD-1/CTLA-4 immune checkpoint blockade treatment, ICBΔT = combined PD-1/CTLA-4 immune checkpoint blockade treatment received after CD4+ T cell depletion. See also supplemental figures 3 and 4 and supplemental tables 7 and 8.

Since the overrepresentation analysis did not include information about the directionality or magnitude of overrepresentation, we wanted to generate a simple quantitative metric that captured both these aspects. To do this, we chose to sum the average log2 fold changes reported by Seurat for each detected gene in the IFN-g response gene set. With this method, a positive value indicates that genes from the gene set skew towards upregulation in the first condition of a given comparison in a given cell type, while a negative value indicates that genes skew towards downregulation. These values indicated that, when treated with ICB treatment, endothelial cells, neutrophils, and luminal cells experienced the most upregulation of IFN-g response genes compared to both the untreated controls and the ICB treated tumors with CD4+ T cell depletion, while macrophages and monocytes appeared to experience the most downregulation (Figure 6B, Supplemental Table 7).

To explore enrichment of up- and downregulated genes more formally, we performed ranked GSEA, using average log2 fold changes as the ranking metric and MSigDB’s hallmark gene sets as the test set (Supplemental Figures 4A-C, Methods). Similar to the results seen with the overrepresentation analysis, we found that the IFN-g response gene set was commonly enriched across multiple cell types. Furthermore, the enrichment results followed similar patterns to those seen using the “sum of fold changes” metric. When looking at the ICB treated condition versus both the control and CD4+ T cell depleted conditions, endothelial cells, neutrophils, and luminal cells showed significant enrichment of upregulated IFN-g response genes, while macrophages and monocytes showed significant enrichment of downregulated genes (Figure 6C, Supplemental Tables 7, 8). This identification of differential responses to IFN-g signaling across cell types again highlights the value of scRNA-seq analysis in examining heterogeneous responses to treatment in individual cell types. Since IFN-g signaling in endothelial cells has been suggested to play multiple roles in the tumor immune response, but its role in ICB treatment response has not been documented, we examined the role of IFN-g signaling in endothelial cells further. ^29, 30^

#### Functional analysis confirms endothelial cells are a principal target of IFN-g and a key mediator of treatment response

To test the role of IFN-g signaling in endothelial cells in response to ICB treatment, we generated a mouse model system where *IFNgR1* could be knocked out specifically in endothelial cells with tamoxifen treatment by crossing CDH5-ERT2-Cre+ mice with *IFNgR1* flox/flox (f/f) mice. Using flow cytometry, we confirmed *IFNgR1* expression was significantly reduced in CD31+ endothelial cells from mice in the knockout conditions compared to intact mice lacking the Cre expressing allele (Supplemental Figure 5). After establishing this model system, we compared tumor growth in *IFNgR1* intact mice with and without ICB treatment to tumor growth in endothelial *IFNgR1* knockout mice with and without ICB treatment (Methods). This comparison revealed that ICB treated knockout mice had tumor growth patterns nearly identical to untreated intact mice, demonstrating that significantly reducing *IFNgR1* expression in endothelial cells negated the anti-tumor effects of ICB treatment (Figure 7A, Supplemental Table 9). Thus, IFN-g response in endothelial cells is necessary for an effective ICB treatment response. Furthermore, untreated tumors in the knockout mice grew more quickly than untreated tumors in the intact mice (Figure 7A). These findings are analogous to previously reported findings which showed that CD4+ T cell depletion in the MCB6C model not only prevented ICB induced tumor rejection, but also led to increased tumor growth even in the absence of ICB treatment, indicating that a basal level of T cell activity restrains tumor growth.^19^ Similarly, the findings presented here indicated that basal levels of IFN-g signaling in endothelial cells restrained tumor growth and upregulation of IFN-g activity in endothelial cells was necessary for tumor rejection upon ICB treatment.

**Figure 7.**
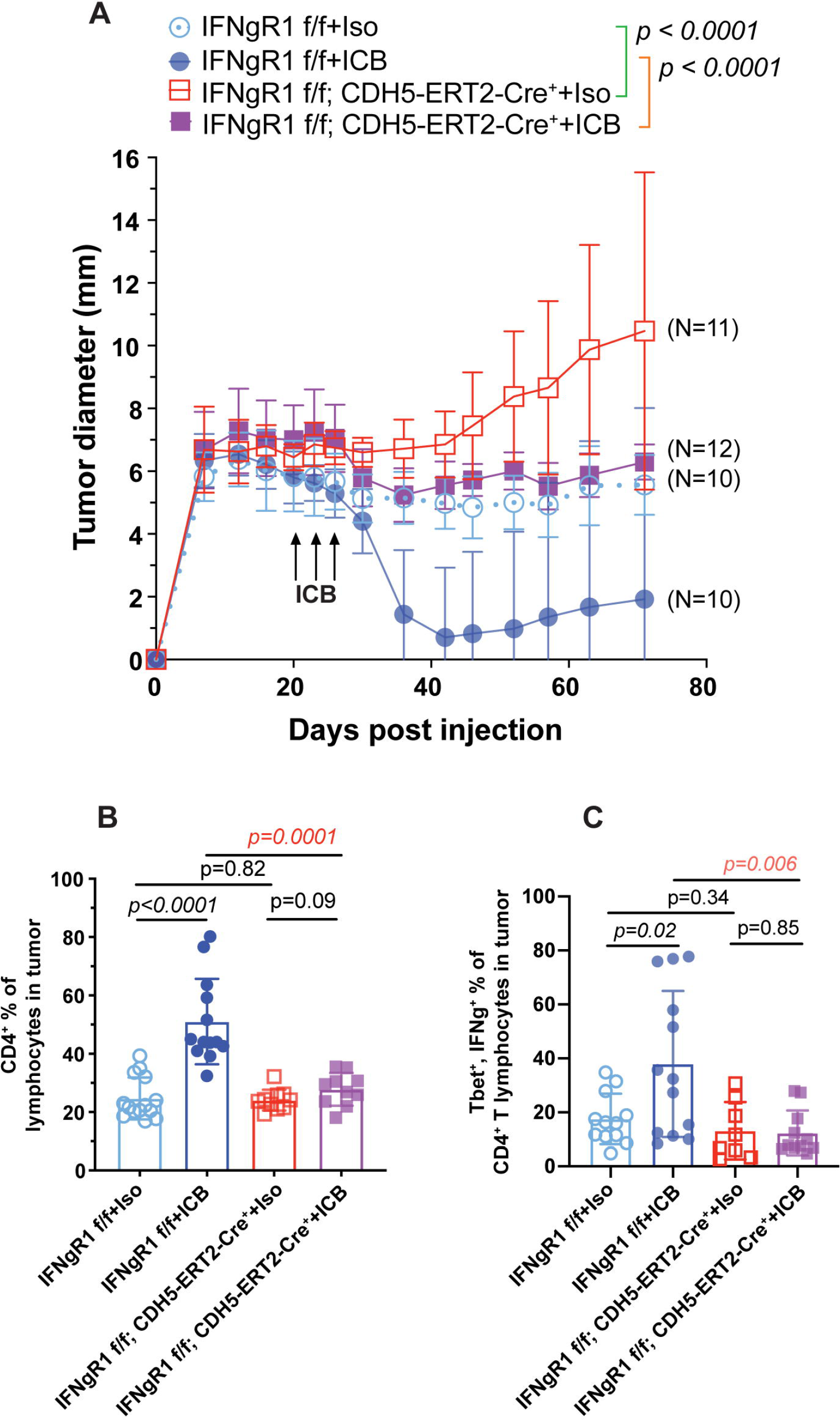
Functional analysis confirms endothelial cells are a principal target of IFN-g and a key mediator of treatment response. (A) Tumor diameter measurements for *IFNgR1* intact and endothelial *IFNgR1* knockout mice with and without ICB treatment over time (pre- and post-treatment). Error bars represent one standard deviation. (B) Bar graphs displaying the percentage of CD4+ lymphocytes in the tumor microenvironment across the same four conditions as (A). Bar height indicates the average percentage across all mice from the given condition. Each point represents the percentage for an individual mouse. Error bars represent one standard deviation. (C) Bar graphs displaying the percentage of *Tbet*+, *IFNg*+ cells detected within the CD4+ T lymphocyte population across the same four conditions as (A). Bar height indicates the average percentage across all mice from the given condition. Each point represents the percentage for an individual mouse. Error bars represent one standard deviation. f/f = flox/flox, ICB = combined PD-1/CTLA-4 immune checkpoint blockade treatment, Iso = rat IgG2a and mouse IgG2b isotype control. See also supplemental figure 5 and supplemental table 9.

Flow cytometric analysis confirmed that ICB treatment induced expansion of CD4+ T lymphocytes in the TME, consistent with previous work.^19^ However, in mice where *IFNgR1* had been knocked out in endothelial cells, this expansion of CD4+ T lymphocytes after treatment was negated (Figure 7B, Supplemental Table 9). Similarly, expansion of *Tbet*+, *IFNg*+ CD4+ T lymphocytes seen after ICB treatment was no longer seen in the knockout condition (Figure 7 C, Supplemental Table 9). Ultimately, these results further indicated that IFN-g signaling in endothelial cells is a key mediator of treatment response and that it underlies expansion of effector T cells in the TME.

## Discussion

To explore mechanisms of response to combined PD-1/CTLA-4 immune checkpoint blockade (ICB) treatment of bladder cancer in individual cell types, we generated scRNA-seq from a mouse model of urothelial carcinoma. The three sample conditions used in this study were untreated tumor, combined PD-1/CTLA-4 ICB treated tumor, and tumor that received combined ICB treatment after CD4+ T cell depletion. In total, we performed scRNA-seq on fifteen samples (five per each condition) and captured over 57,000 cells that passed filtering and were aggregated into a single data set for downstream analysis. Within the aggregated data set, we identified numerous lymphocyte, myeloid, and stromal cell populations. Clustering of the data revealed two distinct epithelial clusters, which we confirmed corresponded to tumor cell populations based on expression of somatic variants and, more specifically, corresponded to distinct basal-like and luminal-like subpopulations based on marker expression.

After identifying cell types present within the aggregated data set, we used differential expression, overrepresentation, and gene set enrichment analysis to explore how individual cell types were responding to treatment. This analysis showed that IFN-g response was commonly perturbed with treatment across multiple cell types, including tumor cells and endothelial cells. Multiple clinical trials exploring human bladder cancer have identified IFN-g pathway activity as being correlated with increased benefit from ICB treatment.^12, 13, 31^ Previous work in the MBC6C model established that IFN-g activity is necessary for ICB treatment response.^19^ While previous research of tumor immunosurveillance models has shown that IFN-g signaling can act through both tumor cell intrinsic and extrinsic mechanisms, the role of IFN-g and its key target cells in ICB treatment response has not been completely defined.^29, 32, 33^

While previous work excluded IFN-g activity in tumor cells as having an essential role in treatment response in the MCB6C model, we had not previously evaluated its role in endothelial cells. Here we establish endothelial cells as a key target of IFN-g activity and further show that loss of IFN-g signaling in endothelial cells impairs expansion of IFN-g producing CD4+ T cells in the TME. Notably, *Cxcl9* and *Cxcl10*, which are mediators of T cell trafficking, were amongst the most upregulated IFN-g response genes in endothelial cells following ICB treatment, suggesting that a key role of IFN-g activity in endothelial cells may be to enable recruitment of T cells to the TME.

We hypothesize a feed-forward model in which ICB treatment induces IFN-g production from CD4+ T cells, which in turn leads to further recruitment of CD4+ T cells to the TME via upregulation of chemoattractant molecules in endothelial cells. However, other roles of IFN-g signaling in endothelial cells could also contribute to treatment response. For example, IFN-g signaling in endothelial cells could induce tumor ischemia or impact vascular permeability, as shown by previous studies.^29, 30, 34^ Ultimately, these results showed that IFN-g response in endothelial cells is a key mediator of treatment response and suggested that strategies which selectively induce IFN-g in endothelial cells in the TME could favorably impact response to ICB treatment as well as other T cell based therapies.

Finally, while these findings support the role of IFN-g signaling in endothelial cells as a key node in treatment response, there are limitations to this analysis. In particular, effective treatment response involves a cascade of events which are still not fully defined. For example, the mechanisms by which T cells in the TME actually kill tumor cells is not elucidated in this system. Ultimately, further analysis will be needed to continue fully characterizing the mechanisms responsible for effective ICB treatment response. Nevertheless, these results underscore the power of scRNA-seq analysis to inform hypotheses that, when coupled with mouse modeling, can help identify cell-type specific signaling nodes that are key to generating an effective immune response.

## Acknowledgements

Malachi Griffith was supported by the National Human Genome Research Institute (NHGRI) of the National Institutes of Health (NIH) under Award Number R00HG007940. Malachi Griffith and Obi Griffith were supported by the NIH National Cancer Institute (NCI) under Award Numbers U01CA209936, U01CA231844, U24CA237719 and U01CA248235. Malachi Griffith was supported by the V Foundation for Cancer Research under Award Number V2018-007. Vivek Arora was supported by the Department of Defense Career Development Award W81XWH-17-1-0562 and Clinical Investigator Award from the Damon Runyon Foundation. We thank Dr. Robert Schreiber at Washington University School of Medicine for gifting us C57BL/6N-*Ifngr1^tm^*^1^*^.1Rds^*/J (IFNgR1^flox/flox^) mice.

## Author contributions

Conceptualization V.K.A., M.G., and O.L.G.; Methodology T.H.P.C. and J.K.B.; Formal Analysis S.L.F., B.F., H.S., M.M., and Z.L.S.; Investigation T.H.P.C.; Writing - Original Draft S.L.F.; Writing - Review and Editing V.K.A., M.G., O.L.G., and S.L.F.; Supervision V.K.A., M.G., and O.L.G.; Funding Acquisition V.K.A., M.G., and O.L.G.

## Declaration of interests

V.K.A. currently serves as an employee of Bristol Myers Squibb and has stock options in the company.

J.K.B currently serves as an employee of Pfizer Inc.

## Methods

### Bulk DNA sequencing, alignment, and variant calling

Whole genome sequencing (WGS) libraries were constructed from genomic DNA isolated from an MCB6C cell line sample and a black 6 (B6NTac) matched normal tail sample using Automated Kapa HYPER PCR free preparation kits (catalog #7962371001 – KK8505) and sequenced on the Illumina NovaSeq 6000 platform. WGS reads were aligned to the GRCm38 reference genome using BWA-MEM. Copy number variant calling was performed using the CNVKit (v0.9.6) batch pipeline.^35^ Whole exome sequencing (WES) libraries were constructed and sequenced similarly to the WGS experiment following hybrid capture selection with the hybrid reagent SureSelect DNA - Mouse All Exon V1 (Agilent). WES reads were aligned to the GRCm38 reference genome using BWA-MEM. Somatic variant calling was performed using common workflow language pipelines provided by the McDonnell Genome Institute (https://github.com/genome/analysis-workflows). Somatic variants were called with Pindel, VarScan, Mutect, and Strelka and combined as previously described.^36–40^ Variants were then filtered based on the criteria of being called by at least two variant callers, normal coverage > 30X, tumor coverage > 30X, normal VAF < 5%, and tumor VAF > 5%.

### Identification of possible driver mutations in whole exome bulk DNA

After filtering somatic variants, a subset of possible driver mutation positions was determined by further filtering the set of somatic mutations down to mutations found in genes that have been previously reported to harbor driver mutations in human bladder cancer (https://www.intogen.org/search?cancer=BLCA). Human gene names were converted to homologous mouse gene names using the Mouse Genome Informatics human and mouse homology report with mammalian phenotype IDs (https://www.informatics.jax.org/homology.shtml). Each mutation was manually reviewed against mutations reported in ProteinPaint (https://proteinpaint.stjude.org/), IntOGen, and Cancer Hotspots (https://www.cancerhotspots.org/#/home).

### Mice

5- to 6-week-old black 6 (B6NTac) male mice were purchased from Taconic Biosciences and were housed in a SPF barrier facility under the guidelines of Institutional Animal Care and Use Committee (protocol # 20-0115) at Washington University. All the in vivo experiments were performed one week after mice were delivered to the animal facility.

### Mouse bladder organoid culture for injection

One previously archived frozen vial of singly suspended MCB6C organoid was thawed at least 2 weeks before mouse injection and expanded weekly in culture at least 2 times. For MCB6C organoid culture expansion, growth factor reduced Matrigel was thawed on ice for minimally 1.5 hours. Pelleted MCB6C cells were washed and resuspended in 1 ml of Advanced DMEM/F12+++ medium (Advanced DMEM/F12 medium [Gibco, catalog #12634010] supplemented with 1% penicillin/streptomycin, 1% HEPEs and Glutamax) and cell concentration was determined by automated cell counter. To establish organoid cultures, 50 ul Matrigel tabs with 10,000 cells/tab were generated and plated on 6-well suspension culture plates, 6 tabs per each well. Tabs were incubated at 37C for 15 minutes until Matrigel was hardened, overlaid with mouse bladder organoid medium (MBO medium - Advanced DMEM/F12+++ medium supplemented with EGF, A-83-01, Noggin, R-Spondin, N-Acetly-L-cysteine and Nicotinamide), and returned to the tissue culture incubator. Organoids were replenished with fresh MBO medium every 3-4 days and also one day before mouse injection.

### Mouse injection with MCB6C organoid cells

A single cell suspension of MCB6C organoid was generated by TrypLE Express (Gibco, catalog #12605010) digestion of organoid Matrigel tabs at 37C for 15 minutes. After digestion, pelleted cells were washed and resuspended in PBS to determine cell concentration. After cell concentration was adjusted to 20 million/ml in PBS, organoid cells were mixed with growth factor reduced Matrigel at 1:1 ratio before subcutaneous injection into the left flank of the mouse (1 million/100 ul cells for each mouse). Tumor development was monitored using digital calipers to assess the length, width, and depth of each tumor. For ICB treatment, each mouse was injected intraperitoneally with 250 ug anti-PD1 (BioXcell, catalog #BE0146, clone RMP1-14) and 200 ug anti-CTLA-4 (BioXcell, catalog #BE0164, clone 9D9) at days 9 and 12 after organoid implantation. For isotype controls, each mouse was injected with 250 ug rat IgG2a (BioXcell, catalog #BE0089, clone 2A3) and 200 ug IgG2b (BioXcell, catalog #BE0086, clone MPC-11). For CD4+ T cell depletion, each mouse was injected intraperitoneally with 250 ug anti-CD4 (BioXcell, catalog #BE0003-1, clone GK1.5) at days 0 and 7 after organoid cell injection/inoculation. Rat IgG2b (BioXcell, catalog #BE0090, clone LTF-2) was used as isotype control for anti-CD4.

### Harvesting MCB6C tumors for single cell RNA-seq, BCR-seq, and TCR-seq

Based on 10x Genomics Demonstrated Protocols, 14 days after organoid implantation, tumors were dissected from euthanized mice, cut into small pieces of ∼2-4 mm^3^, and further processed into dead-cell depleted single cell suspensions following manufacturer’s protocol using Tumor Dissociation Kit and MACS Dead Cell removal Kit (Miltenyi Biotec). Briefly, tumor tissue pieces were transferred to gentleMACS C tube containing enzyme mix before loading onto a gentleMACS Octo Dissociator with Heaters for tissue digestion at 37C for 80 minutes. After tissue dissociation was completed, cell suspension was transferred to a new 50 ml conical tube, and supernatant was removed after centrifugation. Cell pellet was resuspended in RPMI 1640 medium, filtered through a prewetted 70-uM cell strainer, pelleted, and resuspended in red cell lysis buffer and incubated on ice for 10 minutes. After adding wash buffer, cell suspension was pelted and resuspended in wash buffer. To remove dead cells, Dead Cell Removal Microbeads were added to resuspend cell pellet (100 ul beads per 10^7 cells) using a wide-bore pipette tip. After incubation for 15 minutes at room temperature, the cell-microbead mixture was applied onto a MS column. Dead cells remained in the column and the effluent represented to the live cell fraction. The percentage of viable cells was determined by an automated cell counter. Dead cell removal was repeated if the percentage of viable cells did not reach above 90%. Two rounds of centrifugation/resuspension were performed for two rounds in 1xPBS/0.04% BSA using a wide-bore tip. To submit cell samples for single cell RNA-seq analysis, cell concentration was determined by sampling each cell suspension twice, counting each sampling twice, and adjusting to 1,167 cells/ul. 40 ul of each cell suspension was submitted to Genome Technology Access Center/McDonnell Genome Institute (GTAC/MGI) for single cell RNA-seq analysis using the 5’v2 library kit (10x Genomics catalog #PN-1000263) with BCR and TCR V(D)J enrichment kits (10x Genomics catalog #PN-1000016 and #PN-1000005, respectively). cDNA generation and TCR/BCR enrichment were performed according to the Chromium Single Cell V(D)J Reagent Kits User Guide (CG000086 Rev L). The libraries were sequenced on the S4 300 cycle kit flow (2×151 paired end reads) using the XP workflow as outlined by Illumina. FASTQ outputs were generated.

### Alignment, filtering, and clustering of single cell RNA sequencing

Alignment and gene expression quantification were performed with CellRanger count (v5.0, default parameters). Gene-barcode matrices were then imported into Seurat for filtering cells, QC, clustering, etc.^41^ To filter suspected dying cells, cells were clustered before filtering to identify cells clustering based on high mitochondrial gene expression (i.e. the percentage of UMIs per cell mapping to mitochondrial genes). The cutoff for mitochondrial gene expression was based on the percentage that captured the majority of these cells. A cutoff of 12.5% was used across all replicates. Doublets were filtered based on high UMI expression and CellRanger’s reported doublet rate (0.9% per 1000 cells), with the top 0.9% of cells removed from each condition in each replicate. Cutoffs for filtering cells with low feature detection was done by assigning cell types to each cell using the CellMatch method (as described in a previous publication by Petti, et al), identifying cells that did not have enough features for their cell type to be predicted, and calculating the average number of features detected in those cells.^42^ After filtered cells were removed, gene expression values for each gene in the remaining cells were normalized and scaled and variable genes were selected using Seurat with default settings. Principal component (PC) analysis was then performed using these variable genes (npcs = 20). Clustering of cells was performed using 20 PCs and resolution = 0.7. Finally, dimensionality reduction and visualization were performed using Seurat’s tSNE function. B cell and T cell receptors were assembled and identified using the 10x Genomics CellRanger V(D)J pipeline (v5.0, default parameters).

### Assigning cell types using SingleR

Cell types for each cell were annotated with SingleR using expression profiles from the ImmGen dataset (https://www.immgen.org/).^23, 24^ Cell types were manually simplified to B cell (B), CD4+ T cell (CD4), naive CD4+ T cell (CD4.Naive), naive CD8+ T cell (CD8.Naive), CD8+ effector T cell (CD8.Eff), CD8+ memory T cell (CD8.Mem), dendritic cell, endothelial cell, epithelial cell, fibroblast, macrophage, monocyte, neutrophil, natural killer cell (NK), natural killer T cell (NKT), gamma delta T cell (Tgd), and regulatory T cell (Treg).

### Assigning tumor cell subtypes, reclustering, and assigning differentiation scores

To assign tumor cell subtypes, we classified cells that were expressing any level of *Krt5*, but not *Psca* as basal tumor cells and cells that were expressing *Psca*, but not *Krt5* as luminal tumor cells. Cells that were assigned to the same clusters as basal cells using Seurat’s unsupervised clustering were also classified as basal cells. Similarly, cells that were assigned to the same clusters as luminal cells using Seurat’s unsupervised clustering were classified as luminal cells. After assigning these tumor subtype labels, we separated the basal and luminal cell populations from all other cell populations. We then scaled and normalized gene expression and selected variable genes using Seurat’s default methods. Principal component (PC) analysis was then performed using the variable genes (npcs = 20). Clustering of cells was performed using 20 PCs and resolution = 0.7. Finally, dimensionality reduction and visualization were performed using Seurat’s tSNE function. We then assigned differentiation scores to each cell using CytoTRACE (https://cytotrace.stanford.edu/) and calculated the differentiation score as 1 - the CytoTRACE score.^43^

#### Differential expression analysis, overrepresentation analysis, and gene set enrichment analysis in scRNA-seq

All differential expression analyses were performed using Seurat’s FindMarkers function with the Wilcoxon Rank Sum method. P-values were adjusted using Benjamini-Hochberg multiple test correction. All reference gene sets used for overrepresentation analysis and gene set enrichment analysis (GSEA) were from MSigDB.^44, 45^ For all overrepresentation analysis, results were generated using the enricher function from the clusterProfiler package in R. For comparisons of conditions within each cell type, input gene lists for overrepresentation analysis were generated by taking all genes with adjusted p-value < 0.05 and fold change value > ∼1.2 (i.e. abs(log2FC) > 0.26). For comparison of the full basal-like cluster to the full luminal-like cluster, the input gene list for overrepresentation analysis was generated by taking all genes with adjusted p-value < 0.05 and fold change value > ∼3 (i.e. abs(log2FC) > ∼1.58). For comparison of basal-like ICB treated cells to luminal-like ICB treated cells, the input gene list for overrepresentation analysis was generated by taking all genes with adjusted p-value < 0.05 and fold change value > ∼3 (i.e. abs(log2FC) > ∼1.58) and filtering out DE genes meeting the same criteria when comparing basal-like control cells to luminal-like control cells to generate a list of DE genes unique to the ICB condition. GSEA results were generated using UC San Diego and Broad Institute’s GSEA software to run GSEAPreranked with genes ranked by the average log2 fold changes reported by Seurat.

### Generation of CDH5-ERT2-Cre+, IFNgR1 flox/flox (f/f) mice

C57BL/6-*Tg(Cdh5-cre/ERT2)^1Rha^*mice were originally generated by Dr. Ralf H. Adams and purchased from Taconic Biosciences then bred with C57BL/6N-*Ifngr1^tm^*^1^*^.1Rds^*/J (IFNgR1^flox/flox^) mice that were obtained from Dr. Robert Schreiber at Washington University School of Medicine to generate *Cdh5-cre^ERT^*^2^/IFNgR1^flox/flox^ offspring.

### Postnatal deletion of the IFNgR1 gene in the vascular endothelium by Tamoxifen treatment

Tamoxifen (Alfa Aesar, catalog #J63509) was dissolved in corn oil (MilliporeSigma, catalog #C8267) at the concentration of 20 mg/ml in a 37C shaker overnight one day before the treatment began and kept at 4C during the 5-day treatment.

### In vivo tumor experiments by subcutaneous engraftment of bladder cancer organoids

Tumor experiments were performed following methods established previously with modifications.^19^ To improve the engraftment and growth of the organoid cells on mice, Matrigel with high protein concentration (Corning, catalog #354262) was used instead of growth factor reduced Matrigel (Corning, catalog #356231). After organoids were expanded in culture for > 2 weeks and subsequently harvested by TrypLE Express (Gibco, catalog #12605010) treatment, organoid cells were resuspended in 3:1 PBS/high protein concentration Matrigel (instead of 1:1 PBS/growth factor reduced Matrigel) at 10 million cells/ml. 1 million/100 ul of cell/Matrigel mix was subcutaneously injected into the left flank of the mouse, which was performed one week after the completion of Tamoxifen treatment. Tumor growth was monitored twice a week using digital calipers. The mean of long and short diameters was used for tumor growth curves. For ICB treatment, mice were injected with 250 μg/mouse αPD-1 (BioXcell, catalog #BE0146, clone RMP1-14) and/or 200 μg/mouse αCTLA-4 (BioXcell, catalog #BE0164, clone 9D9) i.p. every 3 days from day 15 to 18 after organoid implantation for short term studies, and from day 15 through day 21 from long term studies. 250 μg/mouse rat IgG2a (BioXcell, catalog #BE0089, clone 2A3) and 200 μg/mouse IgG2b (BioXcell, catalog #BE0086, clone MPC-11) were used as isotype controls.

### Antibodies

The following antibodies were used for flow cytometry: Brilliant Violet 510™ anti-mouse CD45 Antibody (BioLegend, catalog #103137, clone 30-F11), PE/Dazzle™ 594 anti-mouse CD3ε Antibody (BioLegend, catalog #100347, clone 145-2C11), FITC anti-mouse CD4 Antibody (BioLegend, catalog #116003, clone RM4-4), PerCP/Cyanine5.5 anti-mouse CD4 Antibody (BioLegend, catalog #116011, clone RM4-4), PE/Cyanine7 anti-mouse CD8α Antibody (BioLegend, catalog #100721, clone 53-6.7), Alexa Fluor® 700 anti-mouse CD8α Antibody (BioLegend, catalog #100729, clone 53-6.7), Brilliant Violet 421™ anti-mouse CD19 Antibody (BioLegend, catalog #115537, clone 6D5), APC anti-mouse/human CD11b Antibody (BioLegend, catalog #101211, clone M1/70), PE/Cyanine7 anti-mouse CD11c Antibody (BioLegend, catalog # 117317, clone N418), Alexa Fluor® 647 anti-mouse CD326 (Ep-CAM) Antibody (BioLegend, catalog #118211, clone G8.8), PerCP/Cyanine5.5 anti-mouse CD326 (Ep-CAM) Antibody(BioLegend, catalog #118219, clone G8.8), Brilliant Violet 421™ anti-mouse/human CD44 Antibody (BioLegend, catalog #103039, clone IM7), PE anti-mouse CD62L Antibody (BioLegend, catalog #104407, clone MEL-14), PE/Cyanine7 anti-mouse Ly-6C Antibody (BioLegend, catalog #128015, clone HK1.4), PerCP/Cyanine5.5 anti-mouse Ly-6G Antibody (BioLegend, catalog #127615, clone 1A8), PE anti-mouse Siglec-F Antibody (BD Biosciences, catalog #552126, clone E50-2440), APC/Cyanine7 anti-mouse CD335 (NKp46) Antibody (BioLegend, catalog #137645, clone 29A1.4), Alexa Fluor™ 700 anti-Foxp3 Antibody (eBioscience, catalog #56-5773-80, clone FJK-16s), PE/Dazzle™ 594 anti-T-bet Antibody (BioLegend, catalog #644827, clone 4B10), APC/Cyanine7 anti-mouse IFN-γ Antibody (BioLegend, catalog #505849, clone XMG1.2), Alexa Fluor® 647 anti-Ki-67 Antibody (BD Biosciences, catalog #561126 clone B56), Biotin anti-mouse CD119 (IFN-γ R α chain) Antibody (BioLegend, catalog #112803, clone 2E2), PE Streptavidin (BioLegend, catalog #405203)

### Flow cytometry

To determine the cellular composition of the tumor, tumors were isolated, minced into small pieces, and digested for 1 hour in DMEM media (MilliporeSigma, catalog #D5796) containing 100 μg/ml Collagenase type IA (Gibco, catalog #17101015), 100 μg/ml Dispase II (MilliporeSigma, catalog #D4693) and 50 U/ml of DNase I (Worthington Biochemical, catalog #LS002006). Cells were washed in ice-cold PBS with 3% FCS and 2 mM EDTA (FACS buffer) and filtered over 70-μm nylon mesh. After red blood cell lysis with ACK solution (Gibco, catalog #A1049201), cells were stained with a Zombie NIR Fixable Viability kit (BioLegend, catalog #423105) for dead cell exclusion followed by Fc-receptor blocking with purified mouse CD16/32 antibody (BioLegend, catalog #101301, clone 93). After cell surface marker staining with fluorescent-conjugated antibodies, cells were fixed and permeabilized using a Foxp3/transcription factor staining kit (eBioscience, catalog #00-5523-00) and intracellularly stained with fluorescent-conjugated antibodies. Flow cytometric data were acquired by Cytek-upgraded 10-color FACScan cytometers at Washington University Siteman Cancer Center Cell Sorting Core facility and analyzed by FlowJo 10 (TreeStar).

### Statistics

Statistical analyses for *IFNgR1* knockout experiments were performed using Prism 8.3.0 (GraphPad). For all tumor growth curve comparisons, a 2-way ANOVA for repeated measures was used. For all other comparisons, an unpaired Student’s t test was used. All tests were 2-tailed. P-values of less than 0.05 were considered significant.

### Study approval

Mice were handled and housed according to protocols approved by the Washington University School of Medicine Institutional Animal Care and Use Committee (protocol # 20-0115).

## Accession numbers

All raw whole genome, exome, and single-cell RNA-seq have been deposited in the Sequence Read Archive (SRA) and is available under BioProject accession number: PRJNA934380.

## Supplemental information

**Supplemental Table 1. Sequencing metrics for bulk DNA and single cell RNA GEX, BCR, and TCR sequencing**

**Supplemental Table 2. Somatic single nucleotide variant (SNV) and copy number variant (CNV) calling results**

**Supplemental Table 3. Mitochondrial gene expression percentages, UMI counts, and gene feature counts for filtering low quality cells**

**Supplemental Table 4. Cell typing assignments from SingleR with Cd79a, Tek, Epcam, and Col3a1 gene expression values and B cell and T cell receptor expression statuses**

**Supplemental Table 5. Tumor status, epithelial cell typing, Epcam expression level, and tumor subtyping for all filtered cells**

**Supplemental Table 6. CytoTRACE scores, differential expression, and overrepresentation results for comparing basal versus luminal tumor cell populations**

**Supplemental Table 7. Overrepresentation False Discovery Rate (FDR) q-values, sums of fold changes for IFN-g response genes, and GSEAPreranked normalized enrichment scores (NES) for comparing conditions across all cell types**

**Supplemental Table 8. Differential expression results for IFN-g response genes in endothelial cells, comparing conditions pairwise**

**Supplementary Table 9: Tumor diameters and lymphocyte population percentages for IFNgR1 intact and endothelial IFNgR1 knockout mice with and without ICB treatment**

