## Supplemental Figures for "Endothelial cells are a key target of IFN-g during response to combined PD-1/CTLA-4 ICB treatment in a mouse model of bladder cancer"

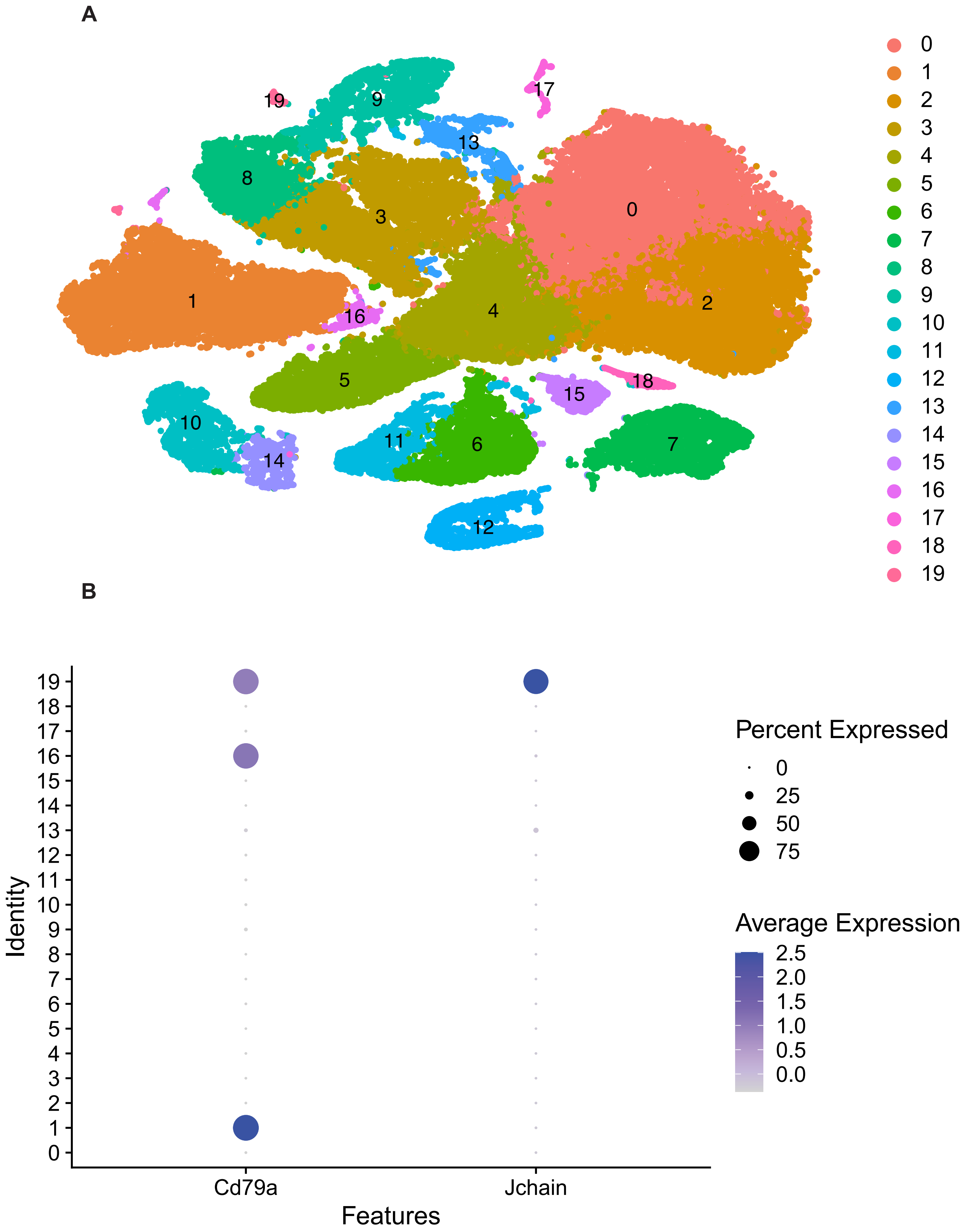

A
B
Percent Expressed
0
25
50
75
Average Expression
2.5
2.0
1.5
1.0
0.5
0.0
Identity
Cd79a
Jchain
Features

**Supplemental Figure 1. tSNE clustering labeled by cluster number, expression of Cd79a and Jchain by cluster number**

(A) tSNE clustering projection of the aggregated data set containing 57,818 cells, across all replicates and conditions, that passed filtering and were not “pruned” by SingleR. Cells are colored by cluster number as assigned by Seurat’s unsupervised clustering algorithm. (B) Dotplot generated in Seurat showing expression of Cd79a and Jchain by cluster number.

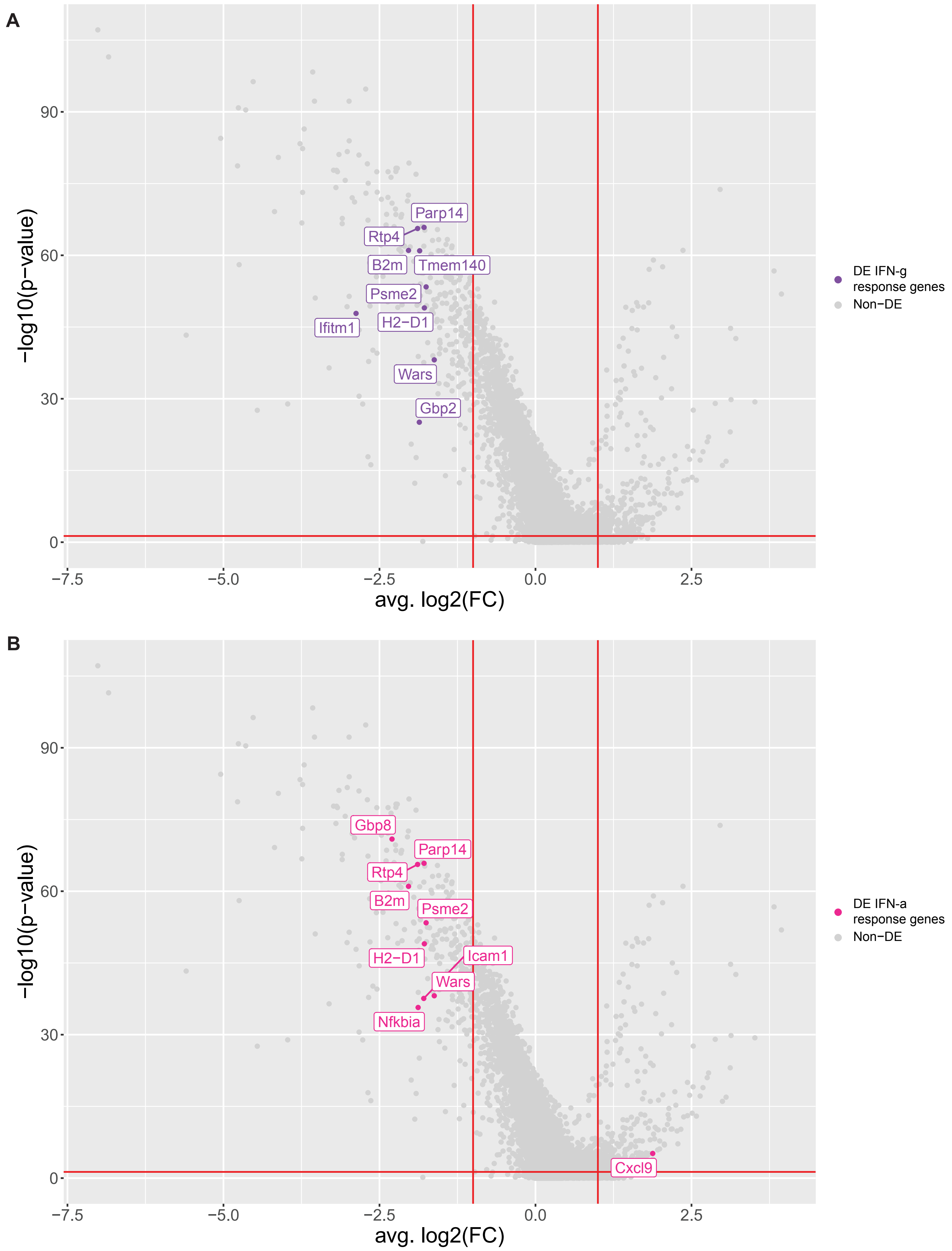

A
Parp14
Rtp4
B2m
Tmem140
Psme2
H2−D1
Ifitm1
Wars
Gbp2
−log10(p−value)
avg. log2(FC)
DE IFN-g response genes
Non−DE
B
Gbp8
Parp14
Rtp4
B2m
Psme2
H2−D1
Icam1
Wars
Nfkbia
Cxcl9
−log10(p−value)
avg. log2(FC)
DE IFN-a response genes
Non−DE

**Supplemental Figure 2. Differentially expressed IFN-g and IFN-a response genes when comparing basal and luminal ICB treated cells**

(A) Volcano plot for differential expression results generated using Seurat's FindMarkers function to compare basal ICB treated cells versus luminal ICB treated cells with IFN-g response genes highlighted in purple. Genes to the left of the leftmost vertical red line are downregulated in basal cells compared to luminal cells. Genes to the right of the rightmost vertical red line are upregulated in basal cells compared to luminal cells. (B) Same volcano plot as (A) with IFN-a genes highlighted in pink. ICB = combined PD-1/CTLA-4 immune checkpoint blockade treatment.

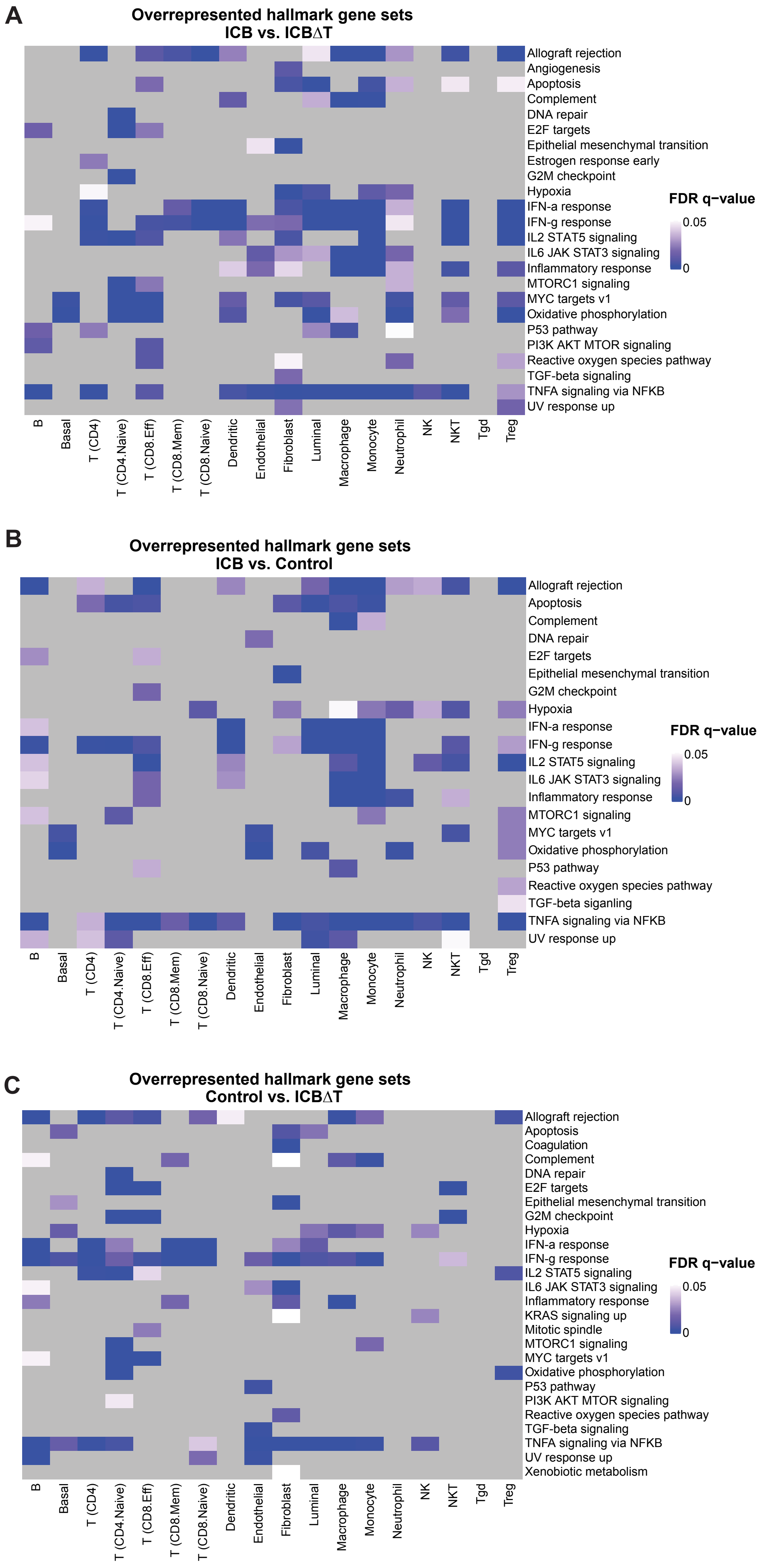
A
Overrepresented hallmark gene sets
ICB vs. ICBΔT
Allograft rejection
Angiogenesis
Apoptosis
Complement
DNA repair
E2F targets
Epithelial mesenchymal transition
Estrogen response early
G2M checkpoint
Hypoxia
IFN-a response
IFN-g response
IL2 STAT5 signaling
IL6 JAK STAT3 signaling
Inflammatory response
MTORC1 signaling
MYC targets v1
Oxidative phosphorylation
P53 pathway
PI3K AKT MTOR signaling
Reactive oxygen species pathway
TGF-beta signaling
TNFA signaling via NFKB
UV response up
FDR q−value
0.05
0
B
Basal
T (CD4)
T (CD4.Naive)
T (CD8.Eff)
T (CD8.Mem)
T (CD8.Naive)
Dendritic
Endothelial
Fibroblast
Luminal
Macrophage
Monocyte
Neutrophil
NK
NKT
Tgd
Treg
B
Overrepresented hallmark gene sets
ICB vs. Control
Allograft rejection
Apoptosis
Complement
DNA repair
E2F targets
Epithelial mesenchymal transition
G2M checkpoint
Hypoxia
IFN-a response
IFN-g response
IL2 STAT5 signaling
IL6 JAK STAT3 signaling
Inflammatory response
MTORC1 signaling
MYC targets v1
Oxidative phosphorylation
P53 pathway
Reactive oxygen species pathway
TGF-beta siganling
TNFA signaling via NFKB
UV response up
FDR q−value
0.05
0
B
Basal
T (CD4)
T (CD4.Naive)
T (CD8.Eff)
T (CD8.Mem)
T (CD8.Naive)
Dendritic
Endothelial
Fibroblast
Luminal
Macrophage
Monocyte
Neutrophil
NK
NKT
Tgd
Treg
C
Overrepresented hallmark gene sets
Control vs. ICBΔT
Allograft rejection
Apoptosis
Coagulation
Complement
DNA repair
E2F targets
Epithelial mesenchymal transition
G2M checkpoint
Hypoxia
IFN-a response
IFN-g response
IL2 STAT5 signaling
IL6 JAK STAT3 signaling
Inflammatory response
KRAS signaling up
Mitotic spindle
MTORC1 signaling
MYC targets v1
Oxidative phosphorylation
P53 pathway
PI3K AKT MTOR signaling
Reactive oxygen species pathway
TGF-beta signaling
TNFA signaling via NFKB
UV response up
Xenobiotic metabolism
FDR q−value
0.05
0
B
Basal
T (CD4)
T (CD4.Naive)
T (CD8.Eff)
T (CD8.Mem)
T (CD8.Naive)
Dendritic
Endothelial
Fibroblast
Luminal
Macrophage
Monocyte
Neutrophil
NK
NKT
Tgd
Treg

**Supplemental Figure 3. Overrepresentation analysis results for hallmark gene sets in pairwise comparisons of conditions across cell types**

(A) Heatmap of False Discovery Rate (FDR) q-values for all hallmark gene sets (rows) that were significantly overrepresented in at least one cell type (columns) when comparing cells from the treated condition to the treated condition with CD4+ T cell depletion. (B) Heatmap of FDR q-values for all hallmark gene sets (rows) that were significantly overrepresented in at least one cell type (columns) when comparing cells from the treated condition to the untreated condition. (C) Heatmap of FDR q-values for all hallmark gene sets (rows) that were significantly overrepresented in at least one cell type (columns) when comparing cells from the untreated condition to the treated condition with CD4+ T cell depletion. For all heatmaps, gray squares indicate that a given gene set was not significant in a given cell type (i.e. q-value > 0.05). ICB = combined PD-1/CTLA-4 immune checkpoint blockade treatment, ICBΔT = combined PD-1/CTLA-4 immune checkpoint blockade treatment received after CD4+ T cell depletion.

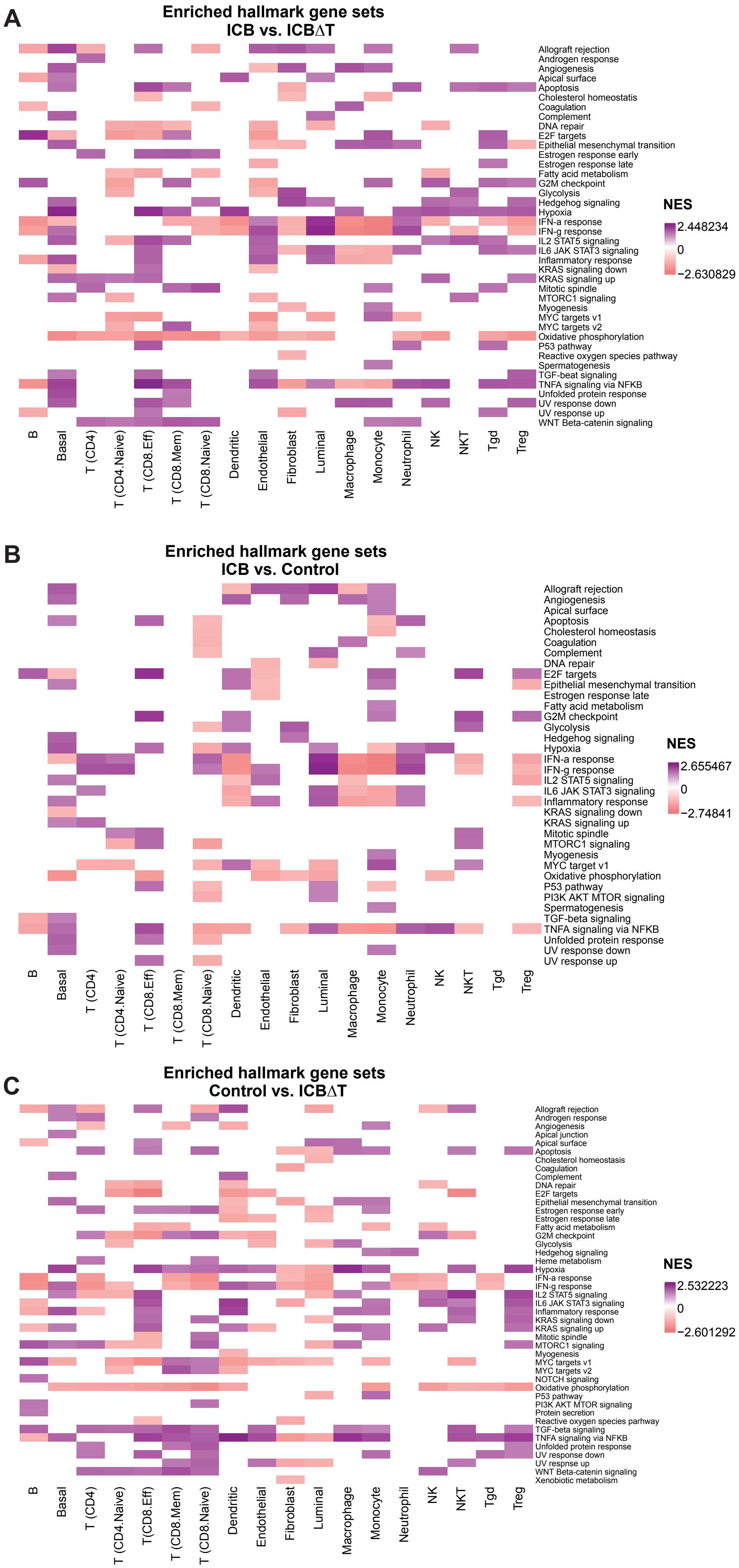

A
Enriched hallmark gene sets
ICB vs. ICBΔT
Allograft rejection
Androgen response
Angiogenesis
Apical surface
Apoptosis
Cholesterol homeostatis
Coagulation
Complement
DNA repair
E2F targets
Epithelial mesenchymal transition
Estrogen response early
Estrogen response late
Fatty acid metabolism
G2M checkpoint
Glycolysis
Hedgehog signaling
Hypoxia
IFN-a response
IFN-g response
IL2 STAT5 signaling
IL6 JAK STAT3 signaling
Inflammatory response
KRAS signaling down
KRAS signaling up
Mitotic spindle
MTORC1 signaling
Myogenesis
MYC targets v1
MYC targets v2
Oxidative phosphorylation
P53 pathway
Reactive oxygen species pathway
Spermatogenesis
TGF-beat signaling
TNFA signaling via NFKB
Unfolded protein response
UV response down
UV response up
WNT Beta-catenin signaling
NES
2.448234
0
−2.630829
B
Basal
T (CD4)
T (CD4.Naive)
T (CD8.Eff)
T (CD8.Mem)
T (CD8.Naive)
Dendritic
Endothelial
Fibroblast
Luminal
Macrophage
Monocyte
Neutrophil
NK
NKT
Tgd
Treg
B
Enriched hallmark gene sets
ICB vs. Control
Allograft rejection
Angiogenesis
Apical surface
Apoptosis
Cholesterol homeostasis
Coagulation
Complement
DNA repair
E2F targets
Epithelial mesenchymal transition
Estrogen response late
Fatty acid metabolism
G2M checkpoint
Glycolysis
Hedgehog signaling
Hypoxia
IFN-a response
IFN-g response
IL2 STAT5 signaling
IL6 JAK STAT3 signaling
Inflammatory response
KRAS signaling down
KRAS signaling up
Mitotic spindle
MTORC1 signaling
Myogenesis
MYC target v1
Oxidative phosphorylation
P53 pathway
PI3K AKT MTOR signaling
Spermatogenesis
TGF-beta signaling
TNFA signaling via NFKB
Unfolded protein response
UV response down
UV response up
NES
2.655467
0
−2.74841
B
Basal
T (CD4)
T (CD4.Naive)
T (CD8.Eff)
T (CD8.Mem)
T (CD8.Naive)
Dendritic
Endothelial
Fibroblast
Luminal
Macrophage
Monocyte
Neutrophil
NK
NKT
Tgd
Treg
C
Enriched hallmark gene sets
Control vs. ICBΔT
Allograft rejection
Androgen response
Angiogenesis
Apical junction
Apical surface
Apoptosis
Cholesterol homeostasis
Coagulation
Complement
DNA repair
E2F targets
Epithelial mesenchymal transition
Estrogen response early
Estrogen response late
Fatty acid metabolism
G2M checkpoint
Glycolysis
Hedgehog signaling
Heme metabolism
Hypoxia
IFN-a response
IFN-g response
IL2 STAT5 signaling
IL6 JAK STAT3 signaling
Inflammatory response
KRAS signaling down
KRAS signaling up
Mitotic spindle
MTORC1 signaling
Myogenesis
MYC targets v1
MYC targets v2
NOTCH signaling
Oxidative phosphorylation
P53 pathway
PI3K AKT MTOR signaling
Protein secretion
Reactive oxygen species parhway
TGF-beta signaling
TNFA signaling via NFKB
Unfolded protein response
UV response down
UV respnse up
WNT Beta-catenin signaling
Xenobiotic metabolism
NES
2.532223
0
−2.601292
B
Basal
T (CD4)
T (CD4.Naive)
T(CD8.Eff)
T (CD8.Mem)
T (CD8.Naive)
Dendritic
Endothelial
Fibroblast
Luminal
Macrophage
Monocyte
Neutrophil
NK
NKT
Tgd
Treg

**Supplemental Figure 4. Gene set enrichment analysis (GSEA) results for hallmark gene sets in pairwise comparisons of conditions across cell types**

(A) Heatmap of normalized enrichment scores (NES) for all hallmark gene sets (rows) that were significantly enriched, using GSEAPreranked method with average log2 fold changes as the ranking metric, in at least one cell type (columns) when comparing cells from the treated condition to the treated condition with CD4+ T cell depletion. (B) Heatmap of NES for all hallmark gene sets (rows) that were significantly enriched, using GSEAPreranked method with average log2 fold changes as the ranking metric, in at least one cell type (columns) when comparing cells from the treated condition to the untreated condition. (C) Heatmap of NES for all hallmark gene sets (rows) that were significantly enriched, using GSEAPreranked method with average log2 fold changes as the ranking metric, in at least one cell type (columns) when comparing cells from the untreated condition to the treated condition with CD4+ T cell depletion. For all heatmaps, white squares indicate that a given gene set was not significant in a given cell type and we do not report the NES for those cell types and gene sets. ICB = combined PD-1/CTLA-4 immune checkpoint blockade treatment, ICBΔT = combined PD-1/CTLA-4 immune checkpoint blockade treatment received after CD4+ T cell depletion.

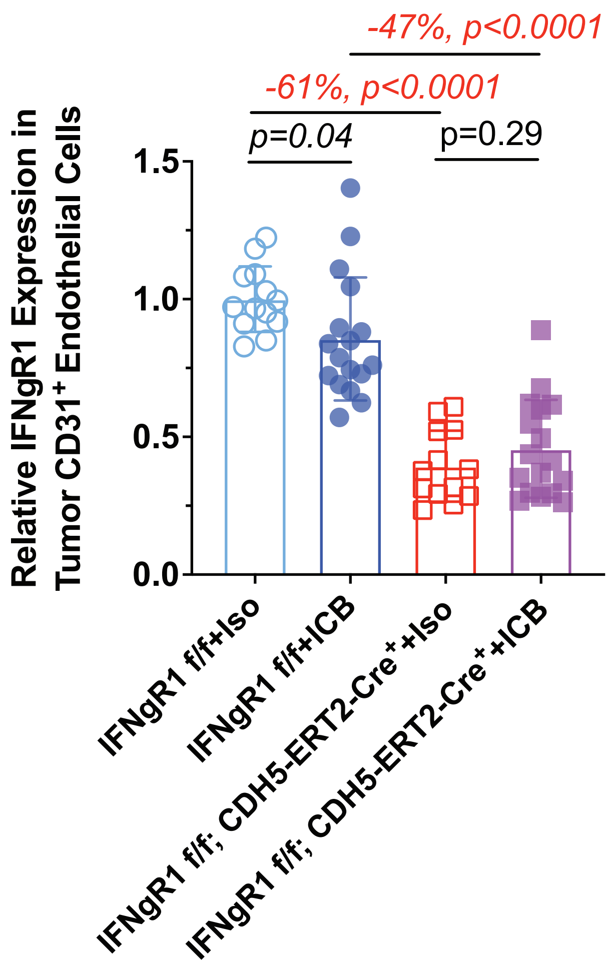

-47%, p<0.0001
-61%, p<0.0001
p=0.04
p=0.29
Relative IFNgR1 Expression in Tumor CD31+ Endothelial Cells
1.5
1.0
0.5
0.0
IFNgR1 f/f+Iso
IFNgR1 f/f+ICB
IFNgR1 f/f; CDH5-ERT2-Cre+ +Iso
IFNgR1 f/f; CDH5-ERT2-Cre+ +ICB

**Supplemental Figure 5. *IFNgR1* expression in *IFNgR1* intact and *IFNgR1* knockout endothelial cells with and without ICB treatment after tamoxifen treatment**
Bar plot comparing the expression of *IFNgR1* in tamoxifen treated IFNgR1 flox/flox mice with or without the CDH5-ERT2-Cre expressing allele which enables endothelial specific Cre expression upon tamoxifen exposure. *IFNgR1* expression was evaluated by flow cytometry in intratumoral endothelial cells identified by expression of CD31. Evaluation was performed in tumors after short term ICB or control treatment. Bar height indicates the average relative expression across all mice from the given condition. Each point represents the relative expression for an individual mouse. Error bars represent one standard deviation. ICB = combined PD-1/CTLA-4 immune checkpoint blockade treatment, Iso = rat IgG2a and mouse IgG2b isotype control.
